# DNA topoisomerase I acts as supercoiling sensor for transcription elongation in *E. coli*

**DOI:** 10.1101/2024.10.07.617030

**Authors:** Vita Vidmar, Céline Borde, Lisa Bruno, Maria Takacs, Claire Batisse, Charlotte Saint-André, Chengjin Zhu, Olivier Espéli, Valérie Lamour, Albert Weixlbaumer

## Abstract

When DNA is transcribed to RNA, the DNA double helix is constantly unwound and rewound to provide access for RNA polymerase (RNAP). This induces DNA supercoiling as a function of transcript length due to over- and under-twisting of the DNA downstream and upstream of RNAP, respectively. Using single-particle cryo-EM and *in vivo* assays we investigated the relationship between bacterial RNAP and DNA Topoisomerase I (TopoI), which removes negative supercoils accumulating upstream of RNAP. TopoI binds to relaxed DNA upstream of RNAP in a manner suggesting a sensory role awaiting the formation of negative supercoils and involving a conformational switch in the functional domains of TopoI. On DNA substrates mimicking negatively supercoiled DNA, TopoI threads one strand into the active site for cleavage while binding the complementary strand with an auxiliary domain. We propose a comprehensive model for DNA relaxation in the context of a transcribing RNAP.

## Introduction

During gene expression, RNA polymerase (RNAP) transcribes the DNA template strand (tDNA) to RNA. The translocation activity of the RNAP elongation complex (EC) causes overwinding of the downstream DNA (positive supercoiling) and underwinding of the upstream DNA (negative supercoiling) as a function of transcript length. This phenomenon, known as the twin supercoiled domain model (Fig. 1a), was initially described by Liu and Wang^1^, and later confirmed both *in vivo* and *in* vitro^2–4^. Locally, DNA supercoiling can reach high levels that might stall transcription by disfavouring forward translocation of RNAP^5,6^.

**Figure 1:**
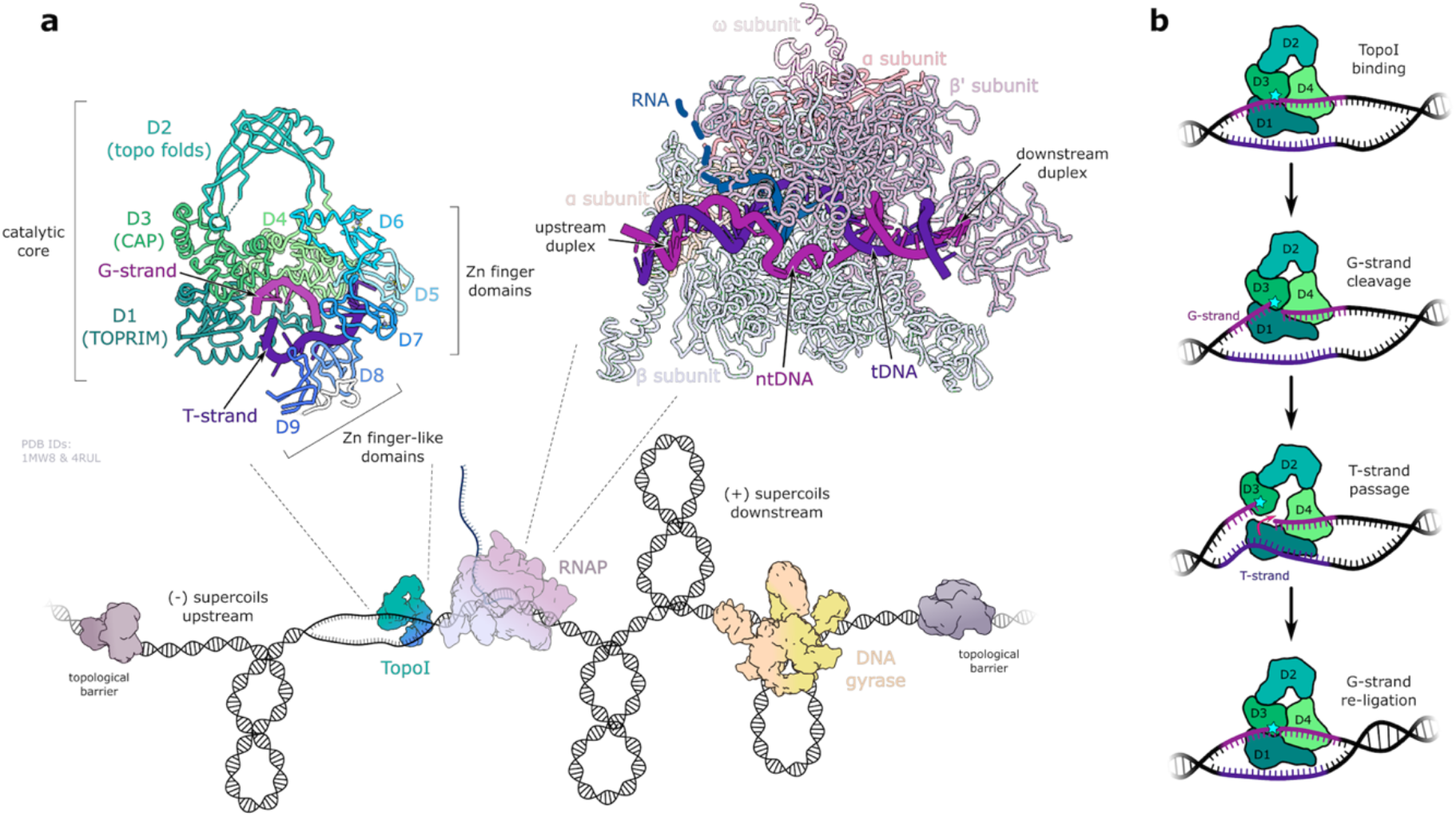
Introduction. **a**, Twin supercoiled domain model with structural details of *E. coli* TopoI (top left) and an RNAP EC (top right). Transcription in combination with topological barriers on DNA induces positive supercoils downstream, which are resolved by DNA gyrase, and negative supercoils upstream, which are relaxed by TopoI. Top left: *E. coli* TopoI composite model (PDB IDs: 1MW8^7^ and 4RUL^9^) consists of a conserved catalytic core (domains D1 – D4) and a modular CTD made of Zn-finger (D5 – D7) and Zn-finger-like domains (D8, D9). Both domains, the catalytic core and the CTD, can bind single-stranded DNA with opposite polarity (G-strand and T-strand). The active site is formed by domains D1 (TOPRIM) and D3 (CAP), the latter having the catalytic Tyr residue. Top right: RNAP consists of five subunits (α_2_ββ’ω). RNA synthesis takes place in the transcription bubble, which is accommodated in the interface between β and β’. The template strand (tDNA) forms a hybrid with the nascent RNA, before reannealing with the non-template strand (ntDNA). **b**, Schematic diagram of DNA relaxation by TopoI. Only the catalytic core (domains D1 – D4) is shown for clarity. The catalytic tyrosine residue in domain D3 is drawn as a cyan star.

To prevent inhibitory excessive DNA supercoiling, transcription is coupled with DNA relaxation. Bacterial topoisomerase I (TopoI), a type IA topoisomerase, relaxes negative supercoils upstream of the EC (Fig. 1a). TopoI acts on single-stranded DNA and relaxes one negative supercoil in each catalytic cycle by passing one DNA strand (T-strand) through a transient break in the complementary strand (G-strand) (Fig. 1b). Crystal structures of the catalytic core or the C-terminal domain (CTD) bound to single-stranded DNA were used to characterize its interaction with DNA^7–9^. However, TopoI preferentially binds the junctions of single-stranded and double-stranded DNA (ssDNA and dsDNA, respectively), a property thought to facilitate the recognition of negatively supercoiled DNA^10,11^. Structural and biochemical data suggested the CTD is required for DNA relaxation^9,11^. Two competing models have been proposed based on structural and biochemical information^7–9^. According to the single-chain model both domains bind the same DNA strand, while in the double-chain model the catalytic core and the CTD bind two complementary strands. DNAse I protection assays provided support for the latter^9^.

In the context of the twin supercoiled domain model, the negatively supercoiled DNA upstream of RNAP is believed to represent a hotspot for TopoI activity^3^. Moreover, in several species, including *E. coli*^12^, *Streptococcus pneumoniae*^13^, *Mycobacterium tuberculosis*, and *Mycobacterium smegmatis*^14^, RNAP and TopoI were proposed to be in direct contact. In *E. coli*, RNAP and TopoI co-purify from cell lysates^12,15^. Direct interaction is supported by genomic data^15^, and one proteomic study^16^, but not another^17^. Binding experiments^12,18^ suggested the RNAP β’ subunit interacts with the C-terminal Zn-ribbon domains of TopoI, which are important for viability^12,15^.

However, the molecular details of the coupling between transcription and DNA relaxation remain unknown. To understand how TopoI operates in the context of the EC, whether its activity is modulated by specific protein-protein contacts or if it is only driven by DNA topology, we used nucleic acid scaffolds to mimic different upstream DNA topologies and obtained single-particle cryo-EM structures of the *E. coli* RNAP-TopoI complex in different functional states. The structures and *in vivo* assays shed light on RNAP-TopoI coupling on relaxed and negatively supercoiled DNA.

## Results

### RNAP-TopoI complex on relaxed DNA

We first sought to understand how RNAP and TopoI cooperate on relaxed DNA before supercoils accumulate or after DNA relaxation. An EC was reconstituted for cryo-EM on a nucleic acid scaffold imitating a transcription bubble, which forms double-stranded DNA immediately upstream of RNAP to mimic locally relaxed DNA (duplex scaffold). Focused cryo-EM reconstructions allowed us to generate a composite map and refine atomic models of an EC and TopoI (Fig. 2a, Supplementary Fig. 1, Supplementary table 1).

**Figure 2:**
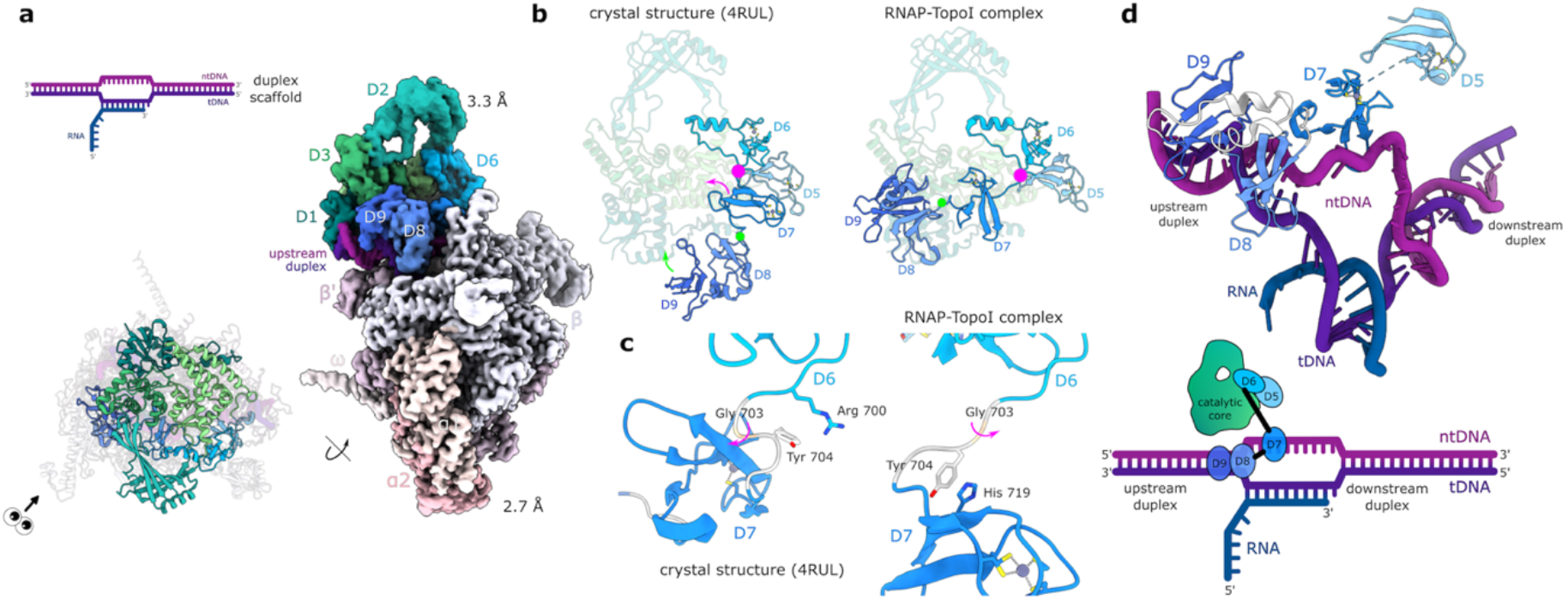
RNAP-TopoI complex on a nucleic acid scaffold mimicking relaxed DNA. **a**, Composite sharpened cryo-EM map and refined atomic model with a schematic representation of the nucleic acid scaffold (top) to show the overall architecture of RNAP-TopoI complex on a duplex scaffold mimicking relaxed DNA. The eye symbol next to the model (RNAP in grey, same orientation in following figures) indicates the viewing direction for the EM map on the right. **b**, Comparison of the conformation of TopoI-CTD in the crystal structure (PDB ID: 4RUL^9^, left) and RNAP-TopoI complex (right). The conformational change consists of two independent rotations around Gly703 in the linker connecting D6 with D7 (magenta circle, 120°), and around residue Lys752 in the linker connecting D7 with D8 (green circle, 50°). **c**, Tyr704 in the TopoI-CTD acts as a conformational switch. In the crystal structure (PDB ID: 4RUL^9^) it interacts with Arg 700 in D6 (left), whereas in the RNAP-TopoI complex, it stacks with His 719 in domain D7 (right). **d**, Binding of the TopoI-CTD to the ssDNA/dsDNA junction of the transcription bubble (D6 which does not bind DNA is omitted for clarity), and schematic representation of TopoI-CTD binding to the DNA fork. D7 binds the single-stranded part of the ntDNA in the transcription bubble in a similar manner as in the crystal structure. D8 is positioned at the junction of single-stranded ntDNA and upstream DNA duplex. D9 is located close to the upstream DNA duplex where the DNA is entirely double-stranded.

TopoI binds DNA at the upstream edge of the transcription bubble in proximity of RNAP, which resembles a canonical EC^19^ (C_α_ RMSD: 0.8 Å). In contrast, TopoI adopts a different conformation compared to the crystal structure of the full-length protein bound to single-stranded DNA^9^. In particular, the TopoI-CTD rotates relative to the catalytic core (Fig. 2b, Supplementary Fig. 2a, Supplementary movie 1) so domains D7 to D9 approach the TopoI body to avoid a steric overlap with RNAP. Tyr704 in the linker connecting domains D6 with D7 can stabilise both, the previously observed and the novel conformation (Fig. 2c, Supplementary movie 1).

This TopoI-CTD conformation aligns DNA-binding residues with the DNA emerging from the EC. In contrast to the TopoI crystal structure^9^, an EC provides ssDNA and dsDNA regions. TopoI domains D7 to D9 bind the edge of the transcription bubble, where the unpaired non-template DNA (ntDNA) reanneals to form the upstream DNA duplex (Fig. 2d, Supplementary Fig. 2b). The DNA geometry, which is different compared to the crystal structure, dictates an alternative TopoI-CTD interaction with the DNA. In the crystal structure, TopoI predominantly contacts DNA bases through aromatic interactions^9^. In the present structure these are often replaced by electrostatic interactions with the DNA backbone (Supplementary Fig. 2b).

In contrast to the TopoI-DNA contacts, the interface of TopoI and RNAP is not well-resolved indicating substantial flexibility. The few protein-protein contacts between RNAP and TopoI involve mainly electrostatic interactions (Supplementary table 2). D5, which was shown to bind DNA, contacts the RNAP β’ subunit instead. Interestingly, in a subset of particles we observe a flexible loop in the TOPRIM domain of TopoI (residues 37 – 60) to form additional contacts with RNAP (Supplementary Fig. 2c, Supplementary table 2).

The surface area buried by protein-protein interactions is ∼900 Å^2^ (TOPRIM loop included). In comparison, TopoI covers ∼1570 Å^2^ of DNA surface, suggesting the latter drives complex stability. To test whether DNA is required for stabilisation, we compared the binding of TopoI to an EC assembled either on the duplex scaffold used for cryo-EM, or a minimal scaffold lacking the transcription bubble and upstream DNA. In electrophoretic mobility shift assays (Supplementary Fig. 2d), an RNAP-TopoI complex formed only with the duplex scaffold and not with the minimal scaffold. Consistently, TopoI co-eluted with RNAP in size-exclusion chromatography only in presence of the duplex scaffold (Supplementary Fig. 2e). We conclude the upstream DNA is required for a stable RNAP-TopoI complex.

### RNAP-TopoI complex on negatively supercoiled DNA

Next, we wanted to understand how TopoI engages negatively supercoiled DNA that accumulates upstream of a transcribing RNAP. To model strand separation arising from high levels of negative supercoiling, we designed nucleic acid scaffolds with mismatched upstream DNA forming a bubble or single-stranded overhangs.

Preliminary cryo-EM reconstructions using scaffolds with short ssDNA overhangs (short-overhang scaffold, Supplementary Fig. 3) confirmed that TopoI-CTD binds the junction of ssDNA and dsDNA, in a manner resembling the duplex scaffold (Fig. 2d). However, the relative orientation of TopoI and RNAP was different, suggesting the TopoI catalytic core binds the non-template strand of negatively supercoiled DNA. This informed two alternative scaffold designs (bubble and long-overhang scaffold) harbouring a consensus TopoI cleavage sequence^15^ in the ntDNA to promote binding to a specific site. DNA cleavage assays confirmed TopoI activity in a complex with RNAP on both scaffolds (Supplementary Fig. 4a, Supplementary Fig. S5a).

**Figure 3:**
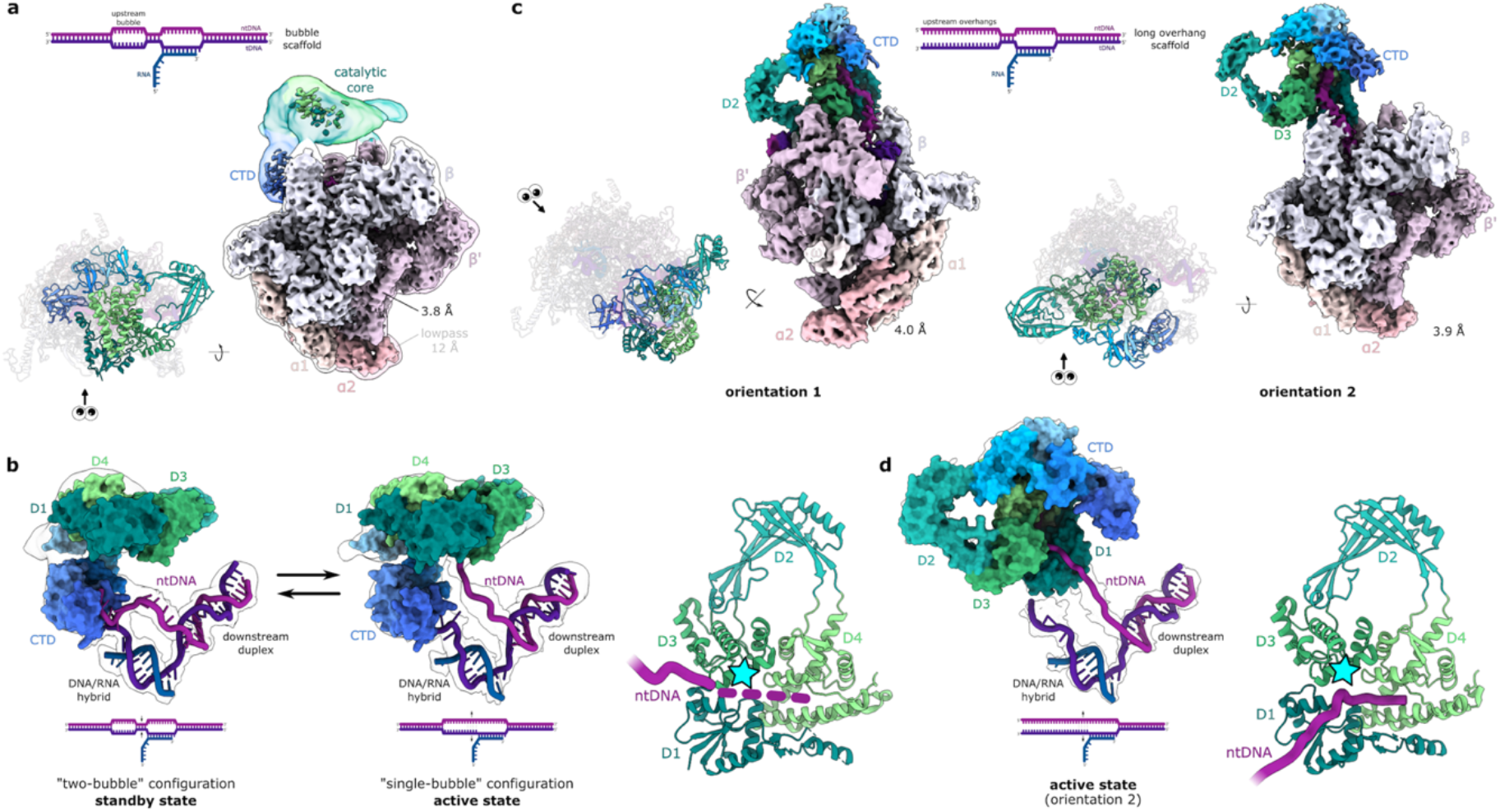
RNAP-TopoI complex on nucleic acid scaffolds mimicking (–) supercoiled DNA. **a**, RNAP-TopoI complex on the bubble scaffold. The EM map in the background is lowpass-filtered to 12 Å. The eye symbol next to the model (RNAP in grey, same orientation in all panels) indicates the viewing direction of the EM map on the right. **b**, Bubble scaffold in a two-bubble (left) and a single-bubble (middle) configuration. The density for the ntDNA in the single-bubble configuration can be traced and leads to the TopoI active site. TopoI (surface) and the scaffold (cartoon) are shown with the EM density (lowpass-filtered to 12 Å) in the background. The TopoI active site Tyr residue in the active configuration (right) is shown as a cyan star. RNAP is omitted for clarity. **c**, RNAP-TopoI complex on the overhang scaffold. In this complex TopoI adopts at least two distinct orientations relative to RNAP. The eye symbols next to the models indicate the viewing direction of the respective EM maps. **d**, The upstream DNA in the overhang scaffold is completely single-stranded; TopoI binds the non-template strand with its active site. TopoI (surface) and the scaffold (cartoon) are shown with the EM density (lowpass-filtered to 12 Å) in the background. RNAP is not shown for clarity.

**Figure 4:**
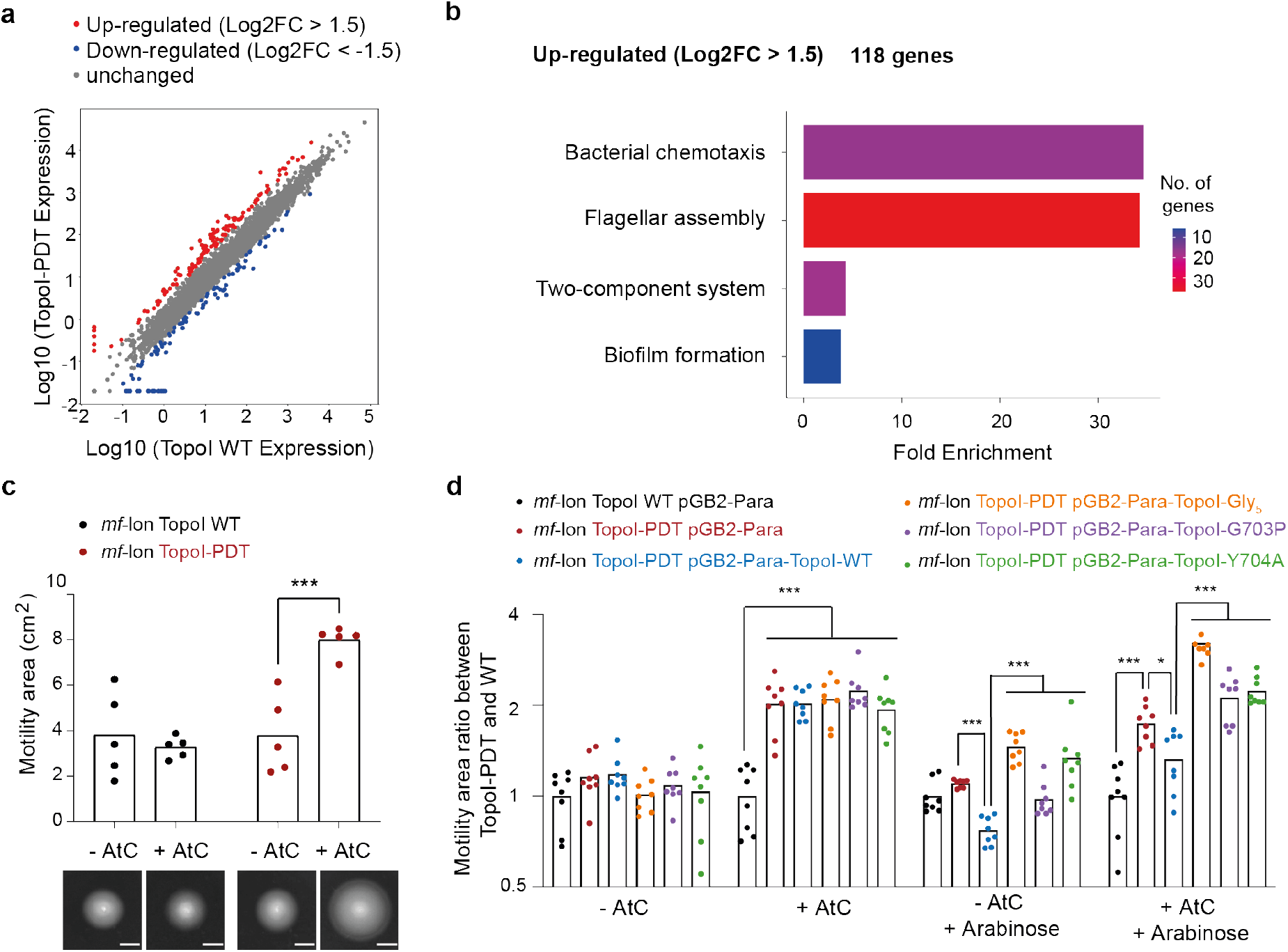
TopoI mutants failed to compensate for TopoI depletion, leading to changes in transcription and up-regulation of flagella. **a**, Scatterplot analysis of differentially expressed genes transcribed under TopoI depletion conditions. A Log2FC ≥ 1.5 threshold was used to assess the significance of differences in gene expression. Red and blue dots represent up- or down-regulated transcripts respectively, and grey dots represent transcripts with no significant changes upon TopoI depletion. **b**, Gene Ontology (GO) enrichment analysis of differentially expressed genes under TopoI depletion conditions using ShinyGo 0.80^27^. For up-regulated genes, significant GO enrichment terms are displayed as a function of the fold-enrichment and the bar colour reflects the number of genes. **c**, *E. coli* motility on LB soft agar of *mf*-lon TopoI WT and *mf*-lon TopoI-PDT in the presence or absence of *mf*-Lon inducer (anhydrotetracycline, AtC) (scale bar: 1 cm). **d**, *E. coli* motility on LB soft agar of *mf*-lon TopoI-WT and *mf*-lon TopoI-PDT with arabinose-inducible expression of TopoI wildtype or mutants. Motility areas were normalized based on the mean of the control condition (*mf*-lon TopoI WT pGB2-Para, -AtC condition). The significance of the two-tailed Mann-Whitney test between averages of both conditions is indicated by ns (not significant) or stars (*: <0.032; **: <0.0021; ***: <0.0002).

The consensus cryo-EM reconstruction of the RNAP-TopoI complex on the bubble scaffold refined to a nominal resolution of 3.8 Å (Fig. 3a, Supplementary Fig. 4, Supplementary table 1). Unexpectedly, TopoI exhibited a high degree of flexibility but the map was sufficiently resolved to unambiguously fit the crystal structure. In this complex, TopoI still binds the EC at the upstream edge of the transcription bubble, but in an orientation rotated by ∼150° relative to RNAP compared to the complex on the duplex scaffold (Supplementary movie 2). The TopoI-CTD is now positioned to bind the template DNA (tDNA) exiting from RNAP, consistent with the strand polarity observed in the full-length crystal structure^9^. 3D variability analysis in CryoSPARC^20,21^ identified two populations with similar TopoI and RNAP relative orientations, but different scaffold configurations (Fig. 3b). In one population, the transcription bubble and the upstream bubble merged. The ntDNA path deviates compared to the canonical EC in the same dataset and connects the transcription bubble with the TopoI catalytic core. The density is not sufficiently resolved to see individual DNA bases but it can be traced to the active site near the junction of TOPRIM and CAP domains. Consistent with the DNA cleavage assay (Supplementary Fig. 4a), modelling of the ntDNA strand suggests the TopoI consensus cleavage sequence can reach the active site in this configuration. We propose this reconstruction represents a state of the complex poised for cleavage and re-ligation. In the second population, the ntDNA emerges from the transcription bubble and re-anneals at the upstream edge forming two bubbles as intended by design. In this conformation, the ntDNA does not enter the TopoI active site suggesting TopoI is in a standby state in contrast to the ‘single-bubble’ conformation.

Unlike on the bubble-scaffold, we observed two distinct TopoI orientations relative to RNAP in cryo-EM reconstructions using the long-overhang scaffold, due to increased flexibility of the ntDNA end. They refined to 4 Å and 3.9 Å nominal resolution, respectively. In the first, TopoI adopts a similar orientation as seen on the bubble scaffold (compare Fig. 3a and Fig. 3c, orientation 1; Supplementary movie 2). The second orientation is related to the first by a ∼145° rotation around an axis that aligns with the exiting ntDNA strand (Fig. 3c, Supplementary Fig. 5, Supplementary movie 2). The TopoI density is better resolved in both reconstructions compared to the bubble scaffold, except for the TopoI-CTD. The CTD is highly mobile causing fragmented density for domains D7 to D9. In both orientations, the ntDNA strand emerging from the transcription bubble extends to the TopoI active site (Fig. 3d), consistent with the active, ‘single-bubble’ conformation on the bubble scaffold, and with the results of the DNA cleavage assay (Fig. 3b, and Supplementary Fig. 5a). Protein-protein contacts between RNAP and TopoI differ in each relative orientation (Supplementary table 2), consistent with our biochemical results suggesting a shared DNA drives complex stability (Supplementary Fig. 2c) and DNA geometry guides the TopoI orientation. Interestingly, the tip of the RNAP β’ subunit clamp domain can form electrostatic contacts with a charged loop in domain D2 of TopoI (Supplementary table 2) in orientation 1 on the long-overhang scaffold and on the bubble-scaffold. This loop has been proposed to facilitate T-strand passage or regulation of DNA gate opening^9,22^. Furthermore, 3D variability analysis^20,21^ identified a transient TopoI state, with a partially open gate and a gap in the EM density between domains D1 and D3 (Supplementary Fig. 5d, Supplementary movie 3). We conclude that TopoI adopts a range of relative orientations to RNAP on scaffolds that mimic negatively supercoiled DNA and any functionally relevant contacts are transient.

**Figure 5:**
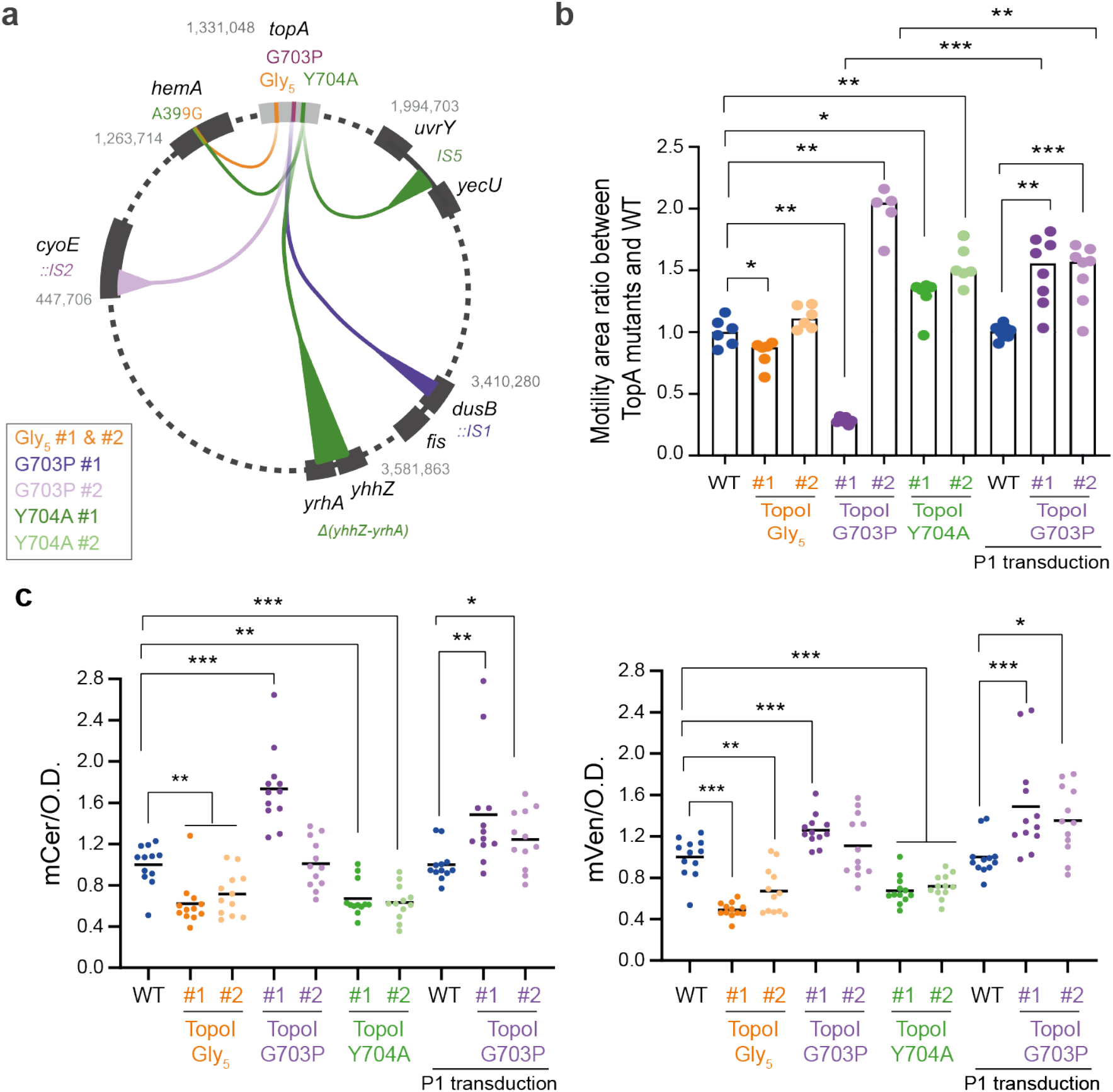
Chromosomal insertion of TopoI mutants leads to mutations, changes in supercoiling and altered gene expression. **a**, Chromosomal mutations acquired in each of two clones (#) of *E. coli* MG1655 TopoI-Gly_5_, TopoI-G703P and TopoI-Y704A. The genome position and affected loci are indicated, except for TopoI-Y704A clone 2, which acquired no additional mutations. Affected loci: *dusB*, tRNA-dihydrouridine synthase B; *hemA*, glutamyl-tRNA reductase; *cyoE* heme O synthase; *uvrY/yecU*, UvrY transcriptional regulator and YecU, a small protein of unknown function; *yhhZ/yrhA*, conserved proteins of unknown function. **b**, *E. coli* motility behaviour on LB soft agar plates of mutant TopoI clones and transductants. Motility areas of mutants are compared to wildtype. **c**, Measurements of reporter gene fluorescence in TopoI mutants. Each point represents an individual well. The fluorescence values were normalized to the wildtype strain. Mutants are compared to wildtype by performing a two-tailed Mann-Whitney test between averages of both conditions and indicated as ns (not significant) or stars (*: <0.032; **: <0.0021; ***: <0.0002).

### *In vitro* functional analysis of TopoI mutants

To investigate the impact of the TopoI-CTD conformational change, we introduced two point mutations. Firstly, we reduced TopoI-CTD flexibility by mutating the pivotal residue, Gly703, to proline (TopoI-G703P). Secondly, we mutated Tyr704 to alanine (TopoI-Y704A) to diminish the stabilising interactions of the two alternative TopoI-CTD conformations (Fig. 2c). Our cryo-EM reconstructions of the RNAP-TopoI complex on negatively supercoiled DNA indicate a potential contact of a charged loop in D2 of TopoI with RNAP (Supplementary table 2). This loop in type IA topoisomerases has been hypothesised to play a role in T-strand passage or gate opening^9,22^. To elucidate its function, we replaced the loop (residues 442 – 448) with five glycines (TopoI-Gly_5_).

We compared the activity of these proteins in DNA relaxation assays *in vitro* (Supplementary Fig. 6a). TopoI-G703P exhibited reduced relaxation activity compared to the wild type, while TopoI-Y704A and TopoI-Gly_5_ showed no discernible effect. This suggests CTD flexibility is necessary for efficient DNA relaxation but the conformational stabilisation observed in the crystal structure and our cryo-EM reconstruction is not required. Likewise, the D2 loop does not appear to be important for DNA relaxation by the isolated enzyme. Next, we tested how CTD mutations (G703P and Y704A) affect binding to the EC on the duplex scaffold. Electromobility shift assays and size-exclusion chromatography revealed reduced binding to the EC (Supplementary Fig. 6b,c). This suggests conformational stabilisation of TopoI-CTD is required for stable binding to an EC, aligning with our structural observations.

**Figure 6:**
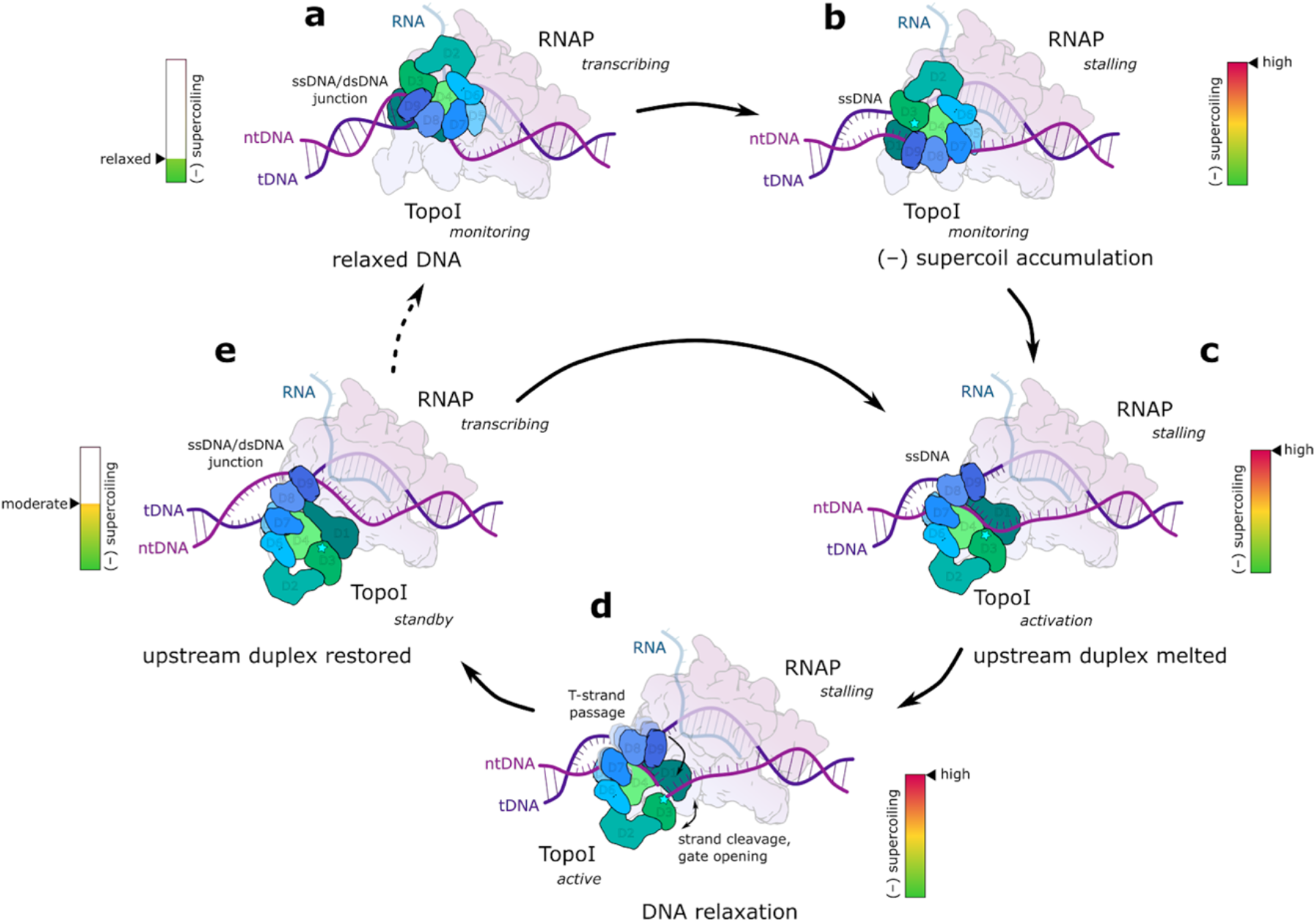
Proposed model for (–) supercoiling detection and relaxation in the context of elongating RNAP. **a**, Detection of DNA topology upstream of an RNAP EC. When DNA is relaxed, the transcription bubble stabilises the preferred DNA geometry for TopoI – the ssDNA/dsDNA junction between the ntDNA strand and the upstream duplex. TopoI can bind the junction with the CTD domains D7 – D9, but not D5. **b**, Melting of the duplex due to excessive (–) supercoil accumulation would change the DNA binding mode of TopoI to entirely single-stranded DNA and potentially allow domain D5 to bind as well, as observed previously in the crystal structure (PDB ID: 4RUL^9^). This transition would necessarily result in the conformational change of the TopoI-CTD, since the conformation in the crystal structure is not compatible with the RNAP EC. **c**, The expansion of the transcription bubble due to melting of the upstream DNA would allow TopoI to access single-stranded DNA with its active site to relax supercoils. **d**, TopoI binds and cleaves the ntDNA strand, while the tDNA strand is bound by its CTD. The direction of CTD movement towards the catalytic core would likely promote the strand passage. **e**, Supercoil relaxation allows re-annealing of the upstream duplex and transcription to resume. At moderate levels of (–) supercoiling (no strand separation), TopoI is sterically hindered from cleaving the non-template strand in the transcription bubble and thus cannot relax the DNA, but remains available to intervene if the supercoiling level increases again. The active site Tyr in TopoI is labelled with a cyan star.

### *In vivo* functional analysis of TopoI mutants

To investigate these interactions *in vivo*, we assessed the impact of the TopoI mutants (TopoI-Gly_5_, TopoI-Y704A and TopoI-G703P) on *E. coli* viability. A TopoI deletion mutant is not viable or acquires compensatory mutations elsewhere in the genome^23^. Therefore, evaluating phenotypes of TopoI mutants affecting the relationship with RNAP is challenging and we engineered an *E. coli* strain encoding a degradable TopoI using the *Mesoplasma florum* Lon protease and its cognate signal peptide PDT (TopoI-PDT, see Methods)^24^. Remarkably, this strain remained viable with no growth defect or change in physiology, despite TopoI being reduced to 15% of its original level upon Lon protease induction (Supplementary Fig. 7a-f).

RNA-seq experiments demonstrated that reduced TopoI levels significantly affect expression of 214 genes (118 upregulated and 96 downregulated, Fig. 4a). TopoI expression increased ∼2-fold, while DNA gyrase, TopoIV and TopoIII were expressed at similar levels. Gene Ontology (GO) analysis indicated a significant upregulation of genes involved in bacterial motility and chemotaxis (Fig. 4b). These changes correlated with a robust increase in bacterial motility after growing overnight in TopoI depletion conditions (Fig. 4c). Whole genome sequencing of the TopoI-PDT strain before and after depletion indicated no other mutations were acquired. We conclude that increased motility is a genuine phenotype reflecting reduced levels of functional TopoI.

We expressed the TopoI mutants from a low copy plasmid controlled by arabinose in the TopoI-PDT strain under depletion conditions (Fig. 4d). Induction of wildtype TopoI restored low motility but expression of the mutants did not, suggesting the mutants lack normal TopoI function. Moreover, in presence of wildtype TopoI levels, the expression of the mutants significantly increased the motility suggesting they may have a dominant negative effect.

Next, we introduced the same mutations directly at the *topA* locus using CRISPR-Cas9 (see Methods). These mutants did not exhibit any growth defects and expressed TopoI nearly at wildtype levels (Supplementary Fig. 8b-f). Complete genome sequencing of two clones for each mutant revealed that, except for TopoI-Y704A clone 2, each clone acquired various additional mutations at the *dusB, hemA, cyoE, uvrY-yecU* and *yhhZ-yrhA* loci (Fig. 5a). The effect of these mutations on DNA topology is unknown, with the exception of the *dusB* mutation which impacts the nucleoid-associated protein Fis, and consequently DNA organisation^25^.

First, we evaluated bacterial motility, which reflects *in vivo* TopoI activity. Both Topo-Y704A clones, and TopoI-G703P clone 2 exhibited increased motility (Fig. 5b). Surprisingly, the motility of TopoI-G703P clone 1 decreased, suggesting the *dusB::IS1* mutation influences this phenotype. To verify this, we transduced *E. coli* MG1655 with TopoI-G703P clone 1 or clone 2 *btuR::Cm* to separate it from the impact of IS elements. Amongst eight transductants of the TopoI-G703P clone 1, seven showed increased motilities (Fig. 5b). As a control, six of eight transductants of the TopoI-G703P clone 2 retained high motilities. PCR analysis revealed that none of the transductants acquired the *dusB::IS1* or the *cyoE::IS2* mutations. The increased motility phenotype for TopoI-G703P and TopoI-Y704A, along with the acquisition of compensatory mutations confirmed that CTD conformation is critical for optimal TopoI functioning *in vivo*. In contrast, the TopoI-Gly_5_ mutation did not change motility.

Next, we tested how these TopoI mutations affect DNA topology of a reporter plasmid. Consistent with the *in vitro* assay, TopoI-Gly_5_ and TopoI-Y704A clone 2 had no effect (Supplementary Fig. 8a and Supplementary Fig. 6a). In contrast, TopoI-G703P clone 1 shifted the reporter plasmid to more supercoiled forms, while TopoI-G703P clone 2 and TopoI-Y704A clone 1 showed more relaxed forms (Supplementary Fig. 8a). However, except for TopoI-Y704A clone 2, the presence of acquired mutations prevents us from drawing strong conclusions.

Finally, we evaluated if TopoI mutants affect RNA polymerase activity by monitoring gene expression using mVenus and mCerrulean fluorescent reporters. To exclude promoter specific or transcription factor related effects, we used λ pR (for mVenus) and the synthetic apFAB67 (mCerulean) promoters^26^. The TopoI-Gly_5_ and TopoI-Y704A mutants had reduced expression of mVenus and mCerrulean. Importantly, TopoI-Y704A clone 2 did not contain additional mutations and we conclude reduced reporter expression is a direct effect of the TopoI-Y704A mutation (Fig. 5c). The reporter expression of the TopoI-G703P mutants differed from each other. Clone 1 containing the *dusB::IS1* mutation showed increased expression, while clone 2 containing the *cyoE::IS2* mutation did not show changes in expression. The Topo-G703P tagged mutation of clone 1 and 2 was transduced in *E. coli* MG1655 and reporter expression was tested (Fig. 5c). Both transduced clones conveyed an increase in reporter expression. Although we cannot exclude the emergence of new mutations, we propose that TopoI-G703P is favorable for gene expression in this context.

Altogether, these results showed that the TopoI mutants affect motility, gene expression and accumulate adaptive secondary mutations, that likely compensate for reduced TopoI function and cause changes in plasmid supercoiling.

## Discussion

TopoI must be able to monitor the topological state of DNA during transcription and intervene when the level of negative supercoiling inhibits EC progression. In line with this function, our results now show that the interaction between the EC and TopoI depends on the presence of DNA and is modulated by the level of DNA supercoiling.

In the RNAP-TopoI complex on relaxed DNA, TopoI binds exclusively through its CTD to the junction of ssDNA and dsDNA at the upstream edge of the transcription bubble (Fig. 2). In contrast, in the crystal structure TopoI binds exclusively single-stranded DNA^9^. The structurally homologous DNA-binding domains of the TopoI-CTD contain positively charged and aromatic residues that interact with the phosphate backbone or DNA bases, respectively. This dual interaction mode enables TopoI to bind single-stranded DNA and junctions between single- and double-stranded DNA, which may enable recognition of DNA topology. In the context of transcription, this distinction allows TopoI to detect when the upstream DNA fails to re-anneal due to excessive negative supercoiling (Fig. 6a,b).

TopoI binding to the RNAP EC on relaxed DNA is accompanied by a rotation of the TopoI-CTD relative to the catalytic core. Its orientation is stabilised by a network of residues that act as a conformational switch (Fig. 2b,c). Moreover, different RNAP-TopoI complexes showed different relative orientations of catalytic core and TopoI-CTD. Domains D5 and D6 are the least flexible and remain associated with the catalytic core. In contrast, domains D7 and D8/D9 move as two independent modules, owing to two hinge points on each side of domain D7. The TopoI-CTD flexibility is important for the interaction with the EC as well as for its activity *in vitro* and *in vivo* because both are impaired in the TopoI-G703P mutant. This is consistent with the proposed function of the TopoI-CTD during DNA relaxation that involves passage of the T-strand^9,11,28^, which would likely require significant movement. However, TopoI-G703P was the only mutant that led to increased reporter gene expression (Fig. 5c), suggesting that downregulation of TopoI activity compared to the wildtype may be beneficial for gene expression. The conformational stabilisation by Tyr704, on the other hand, does not seem to be important for DNA relaxation, but contributes to the EC binding. The presence of either mutation in the CTD (G703P or Y704A) in the chromosomal locus is detrimental, leading to additional adaptive mutations. Surprisingly, we did not observe the well characterized TopoI suppressive mutations leading to gyrase down regulation or TopoIV overexpression^29,30^. However, mutations affecting the *dusB-fis* region were also observed in long-term evolution experiments for TopoI mutants causing high levels of DNA supercoiling^25^. The relationship with supercoiling for the other mutations (*hemA A399G, cyoE::IS2, uvrY-IS5-yecU* and *ΔyhhZ-yrhA*) is unknown. Transduction of these TopoI mutants into the wildtype genetic background causes increased motility phenotypes reflecting lack of normal TopoI function (Fig. 4D, 5B), which may be linked to impairment of its supercoiling sensor function during transcription or hampered EC binding.

In the RNAP-TopoI complex on negatively supercoiled DNA, TopoI adopts an active state and uses the ntDNA as the G-strand (Fig. 3). The ntDNA is threaded from the transcription bubble directly into the active site of TopoI, consistent with our biochemical results. On the other hand, if the upstream DNA reanneals, even if only for a few base pairs, the TopoI active site cannot access the ntDNA strand and thus cannot perform DNA relaxation in the complex with RNAP (Fig. 6c-e). The presence of the active and standby state on negatively supercoiled DNA suggests that TopoI becomes active when supercoiling levels are high enough to cause extensive melting of the upstream DNA, which would likely interfere with RNAP translocation. This agrees with the proposed ‘open’ variation of the twin-supercoiled domain model in *E. coli*, where TopoI in complex with an RNAP EC only partially relaxes the DNA, leaving the residual supercoils to diffuse further upstream^15^ and potentially favour promoter opening for subsequent rounds of transcription.

In addition, TopoI-CTD on the bubble scaffold likely binds the upstream tDNA emerging from RNAP. This supports the previously proposed ‘double-chain’ model of DNA binding, where the TopoI catalytic domain binds the G-strand and TopoI-CTD the T-strand^9^. In the RNAP EC, the G-strand would thus correspond to the ntDNA, and the T-strand to tDNA. However, the mechanism of engaging the TopoI-CTD is likely specific to transcription-coupled regulations in *E. coli*, since the TopoI-CTD is not conserved between species^31^.

Remarkably, we observed a small fraction of particles with a partially opened DNA gate on the long overhang scaffold. Although gate opening does not necessarily depend on DNA cleavage^32,33^, it is an obligatory conformational transition for DNA relaxation. Thus, TopoI in this complex is competent to carry out the complete catalytic cycle of DNA relaxation.

*E. coli* TopoI contains several non-conserved loops, whose function is unclear. Some may play regulatory roles. For example, the charged loop in domain D2 (residues 442 – 448) is thought to be involved in the T-strand passage or regulation of gate opening^9^. Moreover, in two of the RNAP-TopoI complexes on the negatively supercoiled DNA, this loop interacts with the tip of the RNAP clamp domain. A mutant replacing the loop with a Gly_5_ linker (TopoI-Gly_5_) retains the DNA relaxation activity of the wild type but *in vivo* it reduces the expression of the reporter genes (Fig. 5c), suggesting that the TopoI-RNAP interface might play a regulatory role in transcription.

Taken together, we revealed a novel regulatory interplay between RNAP and TopoI, which is beyond simple TopoI recruitment through its CTD as suggested earlier^12,15^. We propose that timely detection of supercoil accumulation behind transcription machinery on one hand and on the other hand tuning of TopoI relaxation activity to ensure optimal supercoiling levels for transcription, have important consequences for proper gene expression and might be vital for the fitness of *E. coli*.

## Data availability

The data that support this study are available from the corresponding authors upon reasonable request. The main cryo-EM reconstructions for RNAP-TopoI on duplex scaffold (composite map: EMD-51259; consensus map: EMD-51252; focused maps: EMD-51253, EMD-51254, EMD-51255, EMD-51256, EMD-51257, EMD-51258), for RNAP-TopoI on bubble scaffold (consensus map: EMD-51260; single-bubble configuration: EMD-51261; two-bubble configuration: EMD-51262), and for RNAP-TopoI on long-overhang scaffold (TopoI orientation 1: EMD-51263; TopoI orientation 2: EMD-51264) reported in this paper were deposited in EM Data Bank under the accession codes listed above.

The atomic models for RNAP-TopoI on duplex scaffold (9GDA), for RNAP-TopoI on bubble scaffold (consensus model: 9GDB; single-bubble configuration: 9GDC; two-bubble configuration: 9GDD), and for RNAP-TopoI on long-overhang scaffold (TopoI orientation 1: 9GDE; TopoI orientation 2: 9GDH) were deposited in the Protein Data Bank under the listed IDs.

Supplementary cryo-EM maps for the RNAP-TopoI on duplex scaffold with TOPRIM loop resolved (composite map: EMD-51267; consensus map: EMD-51265; and focused TopoI map: EMD-51266), for the RNAP-TopoI on long-overhang scaffold (TopoI orientation 3: EMD-51268; and open TopoI gate: EMD-51269) and for RNAP-TopoI on short-overhang scaffold (TopoI orientation 1: EMD-51270; and TopoI orientation 2: EMD-51271) were also deposited in the EM Data Bank.

RNAseq data was deposited in the NCBI GEO repository under accession number GSE273976.

## Acknowledgments

We thank Alexandre Durand and Nils Marechal from the Structural biology platform at the IGBMC, Simon Fromm from the EMBL Heidelberg Imaging Centre, and Alessandro Grinzato from the European Synchrotron Radiation Facility (ESRF) for their help with EM data collection. We acknowledge the ESRF for the provision of microscope time on CM01 and we acknowledge iNext discovery for providing access to the EMBL Heidelberg (VID 26846). The authors would like to acknowledge the High-Performance Computing Center of the University of Strasbourg for supporting this work by providing access to computing resources. We thank Romain Koszul and Justine Groseille for their help with shotgun sequencing data collection. We thank the CIRB imaging facility. Illustrations were generated with BioRender under a personal subscription. This work was supported by European Research Council starting grant TRANSREG 679734 (AW), the Agence Nationale de la Recherche grant ANR-19-CE11-0001-01 (VL), ANR-21-CE11-0040-01 (AW, VL and OE), and ANR-21-CE12-0032 (OE). The authors acknowledge the support and the use of resources of the French Infrastructure for Integrated Structural Biology FRISBI ANR-10-INBS-05 and of Instruct-ERIC. This work of the Interdisciplinary Thematic Institute IMCBio+, as part of the ITI 2021-2028 program of the University of Strasbourg, CNRS and Inserm, was supported by IdEx Unistra (ANR-10-IDEX-0002), and by SFRI-STRAT’US project (ANR-20-SFRI-0012) and EUR IMCBio (ANR-17-EURE-0023) under the framework of the France 2030 Program. VV is a doctoral fellow supported by EUR IMCBio funds and the Ligue Nationale contre le Cancer. CB is a post-doctoral fellow supported by the Agence Nationale de la Recherche ANR-21-CE12-0032 grant.

## Author contributions

V.V. performed biochemical characterization, sample preparation, cryo-EM data collection, data processing, and model refinement. L.B. and C.B. performed in vivo experiments. L.B., M.T. C.S.A., C.B. and C.Z. provided reagents and helped with data processing. All authors contributed to analysis of the data and interpretation of the results. O.E., V.L., and A.W. supervised the studies. V.V., C.B., O.E., V.L., and A.W. wrote the manuscript.

