## Supplementary material for "DNA topoisomerase I acts as supercoiling sensor for transcription elongation in *E. coli*"

**This file includes:**

- Supplementary figures (1 to 8)
- Supplementary tables (1 and 2)
- Supplementary movies (1 to 3)
- Methods
- References

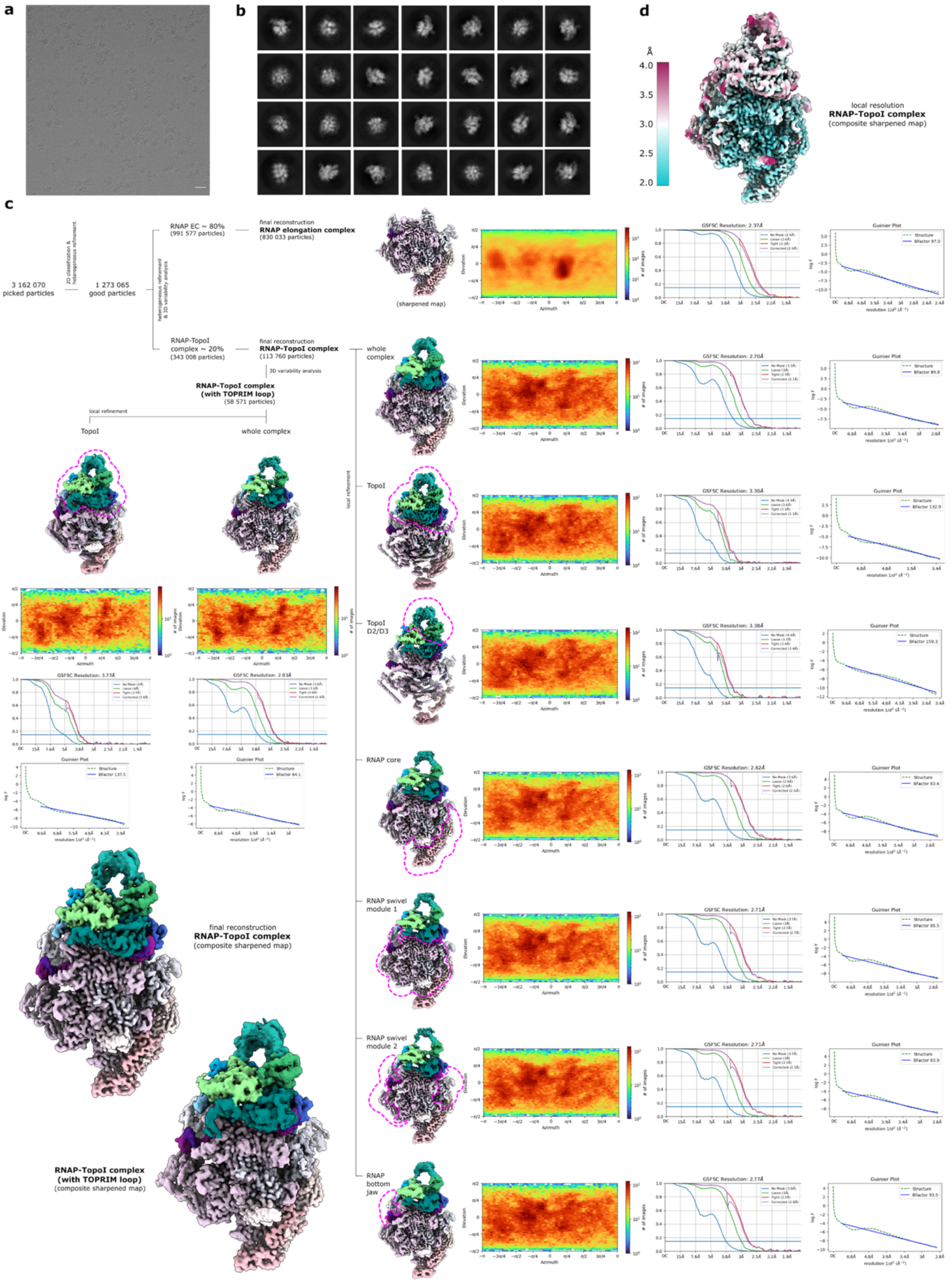

**Supplementary figure 1: Data collection and refinement for RNAP-TopoI complex on duplex scaffold. a**, A representative micrograph. The scale bar is 200 Å. **b**, Representative 2D classes in the dataset. We could not observe classes corresponding to the RNAP-TopoI complex due to its low occupancy, but we could identify them in 3D classification. **c**, Processing and classification tree with the corresponding particle orientation plots, GSFSC curves and Guinier plots of the corresponding final reconstructions. Masked regions in local refinements are labelled with pink dashed line. **d**, Composite sharpened map of RNAP-TopoI complex coloured by the local resolution at FSC threshold 0.143.

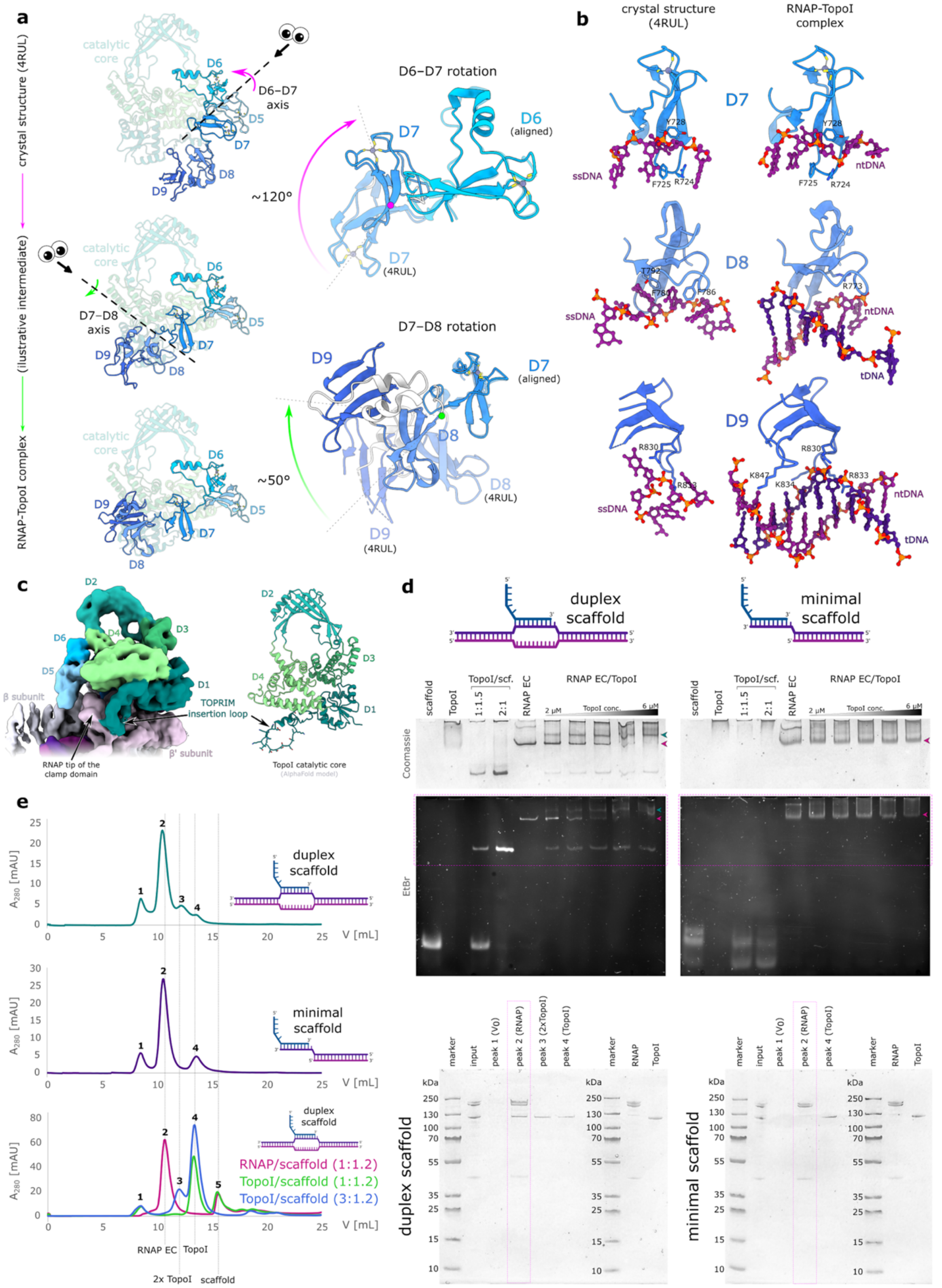

**Supplementary figure 2: RNAP-TopoI complex on duplex scaffold. a**, Comparison of the conformation of TopoI-CTD in the crystal structure (PDB ID: 4RUL[9], top) and RNAP-TopoI complex (bottom) in two views (left vs. right). The conformational change consists of two independent rotations around linkers connecting D6 with D7, and D7 with D8. Domain D7 is rotated around Gly703 by about 120° in the linker connecting D6 with D7. Domains D8 and D9 further rotate relative to D7 around residue Lys752 by about 50° in the linker connecting D7 to D8. The rotation axes are represented with black dashed lines (left) or magenta and green circles (right) and the direction of movement is indicated with arrows. The viewing direction along the rotation axis in the bottom is indicated by eye symbols in the top. **b**, Detailed comparison of DNA binding by TopoI-CTD in the full-length crystal structure (left, PDB ID: 4RUL) and in RNAP-TopoI complex (right). In both crystal structure and RNAP-TopoI complex, domain D7 residues F725, Y728 form $\pi$-$\pi$ stacking with bases. Additionally, residue R724 interacts with a base in the crystal structure but with the phosphate backbone in the RNAP-TopoI complex. R744 forms an extra interaction with the backbone in the RNAP-TopoI complex. D8, which has analogous interactions to D7 in the crystal structure (F780 and F786 that stack with DNA bases), interacts differently in the cryo-EM structure (only R773 interaction with the backbone). D9 in crystal structure interacts with bases as well as the charged DNA backbone (R830, R883), whereas in the complex with transcription bubble it binds the dsDNA, where it forms electrostatic interactions with phosphates (R830, K834) and R833 is inserted into the minor groove. **c**, Map of a particle subset that indicates an interaction of the TOPRIM insertion loop (residues 37 – 60) of TopoI with RNAP. **d**, Electromobility shift assay of RNAP-TopoI complex. An RNAP EC was assembled using either duplex or minimal scaffold (with only a partial transcription bubble and no upstream DNA) and incubated with increasing amounts of TopoI. Complexes corresponding to RNAP EC and RNAP EC-TopoI are highlighted with pink and green arrows in gels stained with Coomassie (top) or ethidium bromide (bottom), respectively. **e**, Size exclusion chromatography and SDS-PAGE of RNAP-TopoI complexes assembled on a duplex scaffold (top) or on a minimal nucleic acid scaffold (middle) or run individually on duplex scaffold (bottom). Duplex scaffold was used as a control, and two different molar ratios of TopoI to scaffold were tested, since two TopoI monomers can bind the scaffold when present in a molar excess (peak 3). TopoI co-elutes with RNAP (peak 2) only in presence of a sufficiently long upstream DNA (compare top vs. middle).

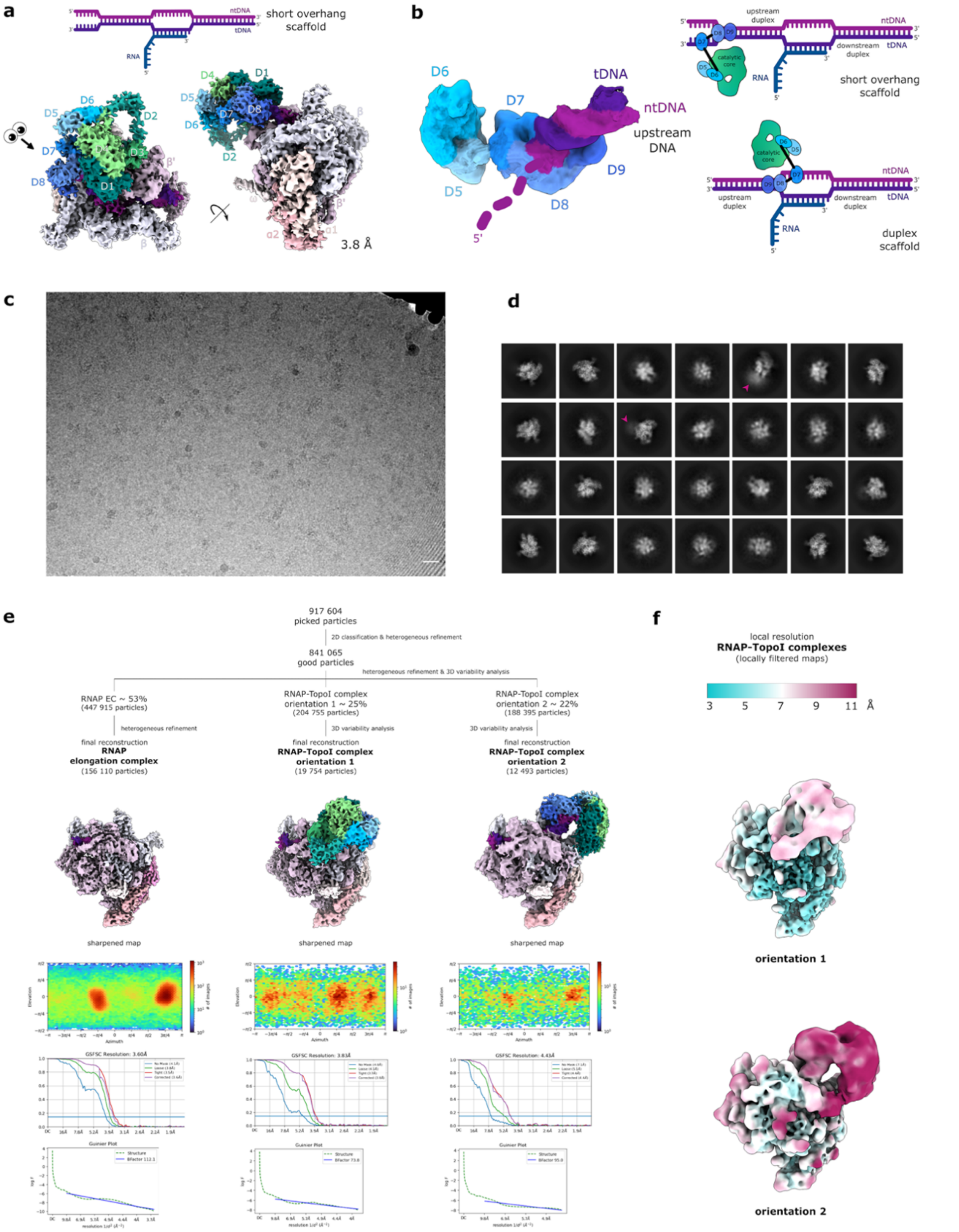

**Supplementary figure 3: RNAP-TopoI complex on short overhang scaffold. a**, Sharpened cryo-EM map with a schematic representation of the nucleic acid scaffold (top). **b**, Binding of the TopoI-CTD to the ssDNA/dsDNA junction on the upstream DNA (left) and a schematic comparing TopoI binding on short overhang scaffold and duplex scaffold (right). A lowpass-filtered (10 Å) map showing TopoI-CTD and the upstream DNA is shown for clarity. The single-stranded ntDNA overhang (not bound by TopoI) is shown as a dashed line. **c**, A representative cryo-EM micrograph. The scale bar is 200 Å. **d**, Representative 2D classes. The extra density for TopoI in some classes is labelled with arrows. **e**, Processing and classification tree with the corresponding particle orientation plots, GSFSC plots and Guinier plots of the corresponding final reconstructions. **f**, Maps of the RNAP-TopoI complex in both conformations filtered and coloured by the local resolution at FSC threshold 0.5. Please note that only maps have been deposited for the RNAP-TopoI complex on short overhang scaffolds with the two TopoI orientations in the EMDB (EMD-51270, EMD-51271, respectively).

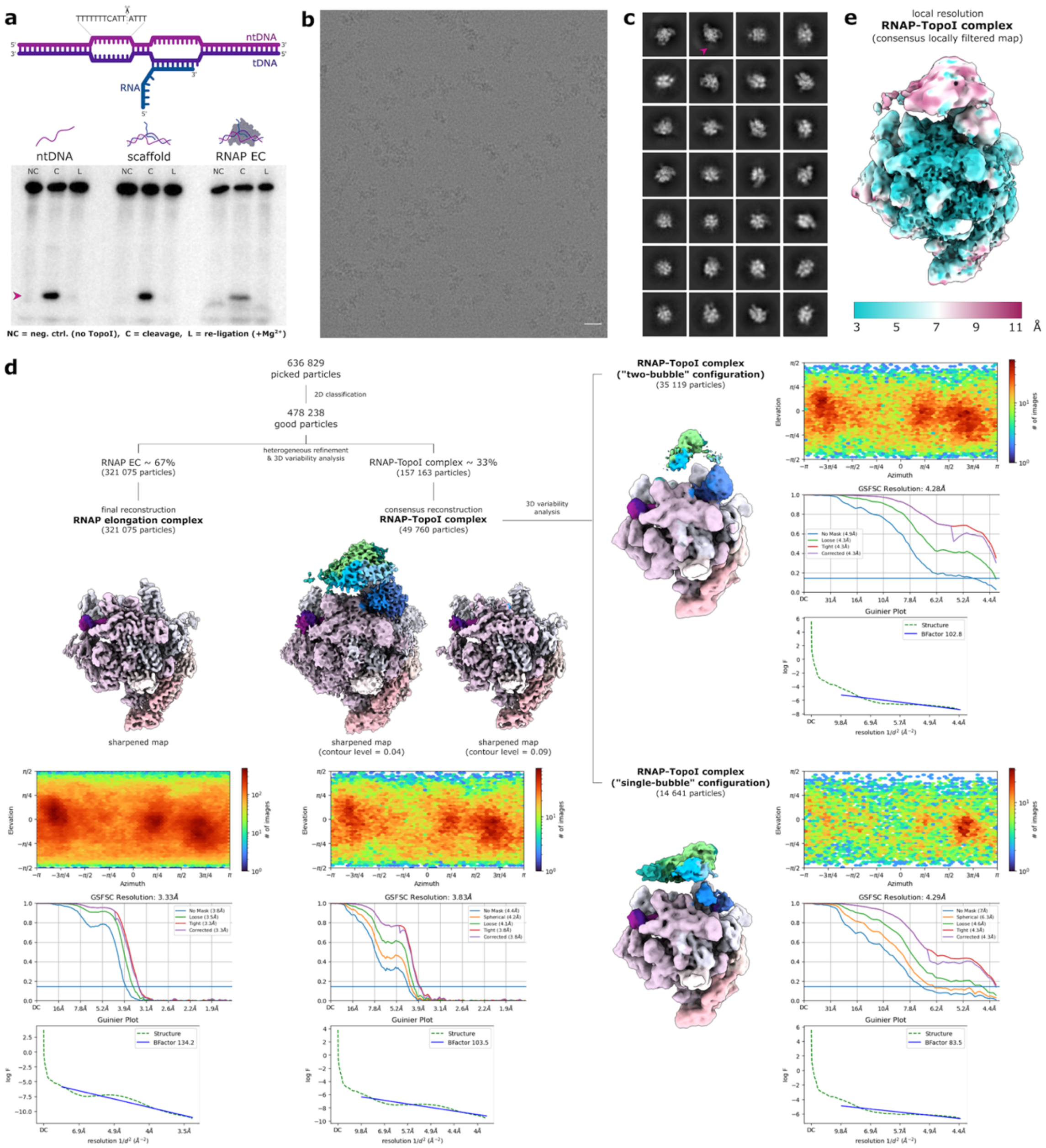

**Supplementary figure 4: Data collection and refinement for RNAP-TopoI complex on bubble scaffold. a**, DNA cleavage assay on bubble scaffold with the consensus TopoI cleavage sequence[15] on the non-template strand (ntDNA). The cleavage intermediate (lanes C), which can be trapped in absence of magnesium, is labelled with a pink arrow and is re-ligated upon addition of magnesium (lanes L, $+Mg^{2+}$). **b**, A representative cryo-EM micrograph. The scale bar is 200 Å. **c**, Representative 2D classes in the dataset. Due to low occupancy of RNAP-TopoI complex and high flexibility of TopoI, the complex was not observed in 2D classes, but could be identified upon 3D classification. **d**, Processing and classification tree with the corresponding particle orientation plots, GSFSC plots and Guinier plots of the corresponding final reconstructions. The consensus map of RNAP-TopoI complex is shown at two different contour levels. **e**, Consensus map of the RNAP-TopoI complex filtered and coloured by the local resolution at FSC threshold 0.143.

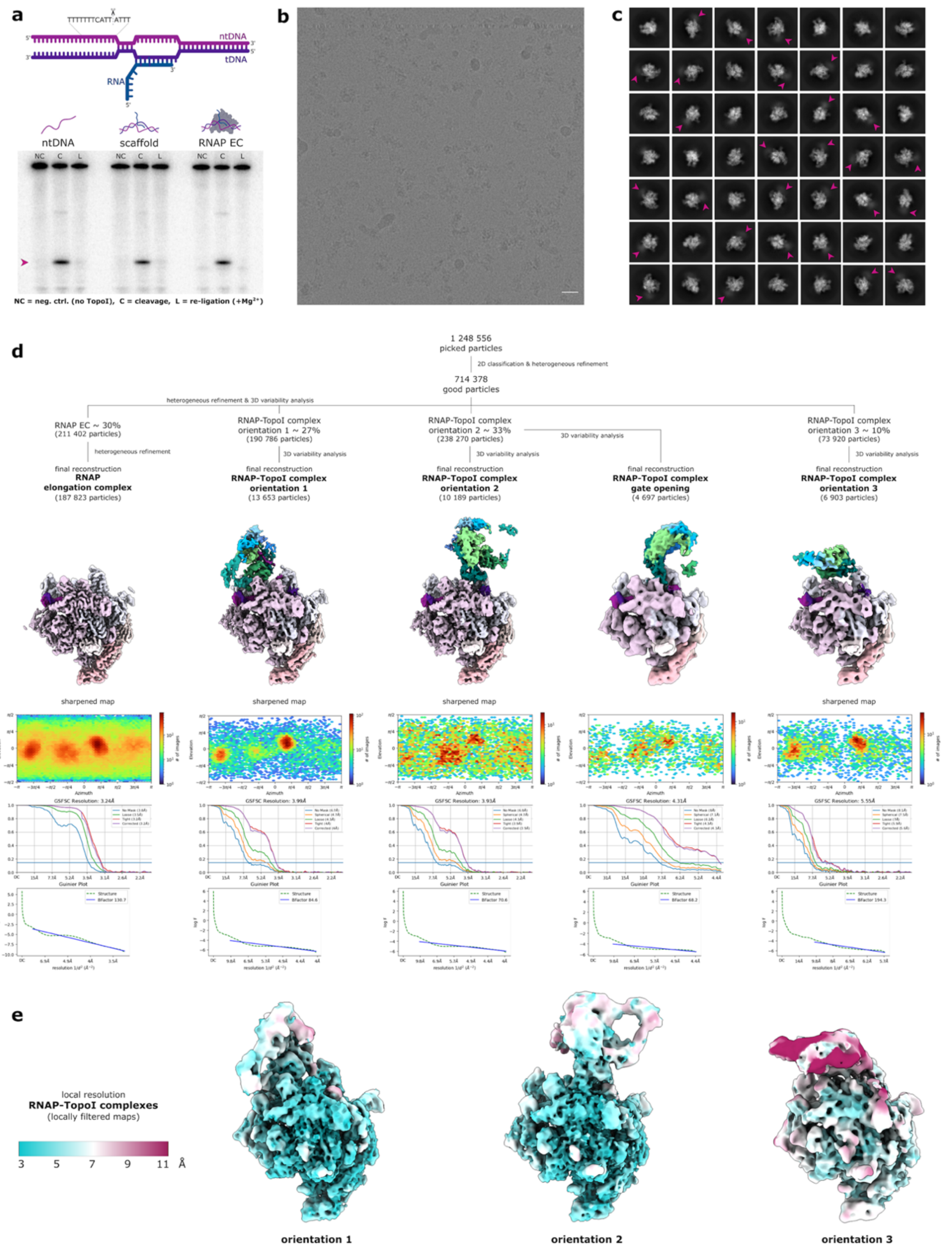

**Supplementary figure 5: Data collection and refinement for RNAP-TopoI complex on long overhang scaffold.** **a**, DNA cleavage assay on long overhang scaffold with the consensus TopoI cleavage sequence[15] on the non-template strand (ntDNA). The cleavage intermediate (lanes C), which can be trapped in absence of magnesium, is labelled with a pink arrow and is re-ligated upon addition of magnesium (lanes L, +$Mg^{2+}$). **b**, A representative micrograph. The scale bar is 200 Å. **c**, Representative 2D classes. The extra density for TopoI in some classes is labelled with arrows. **d**, Processing and classification tree with the corresponding particle orientation plots, GSFSC curves and Guinier plots of the corresponding final reconstructions. Please note that for orientation 3 and for the open TopoI-gate, only maps have been deposited in the EMDB (EMD-51268, and EMD-51269, respectively). **e**, Maps of the RNAP-TopoI complex in all three conformations filtered and coloured by the local resolution at FSC threshold 0.5.

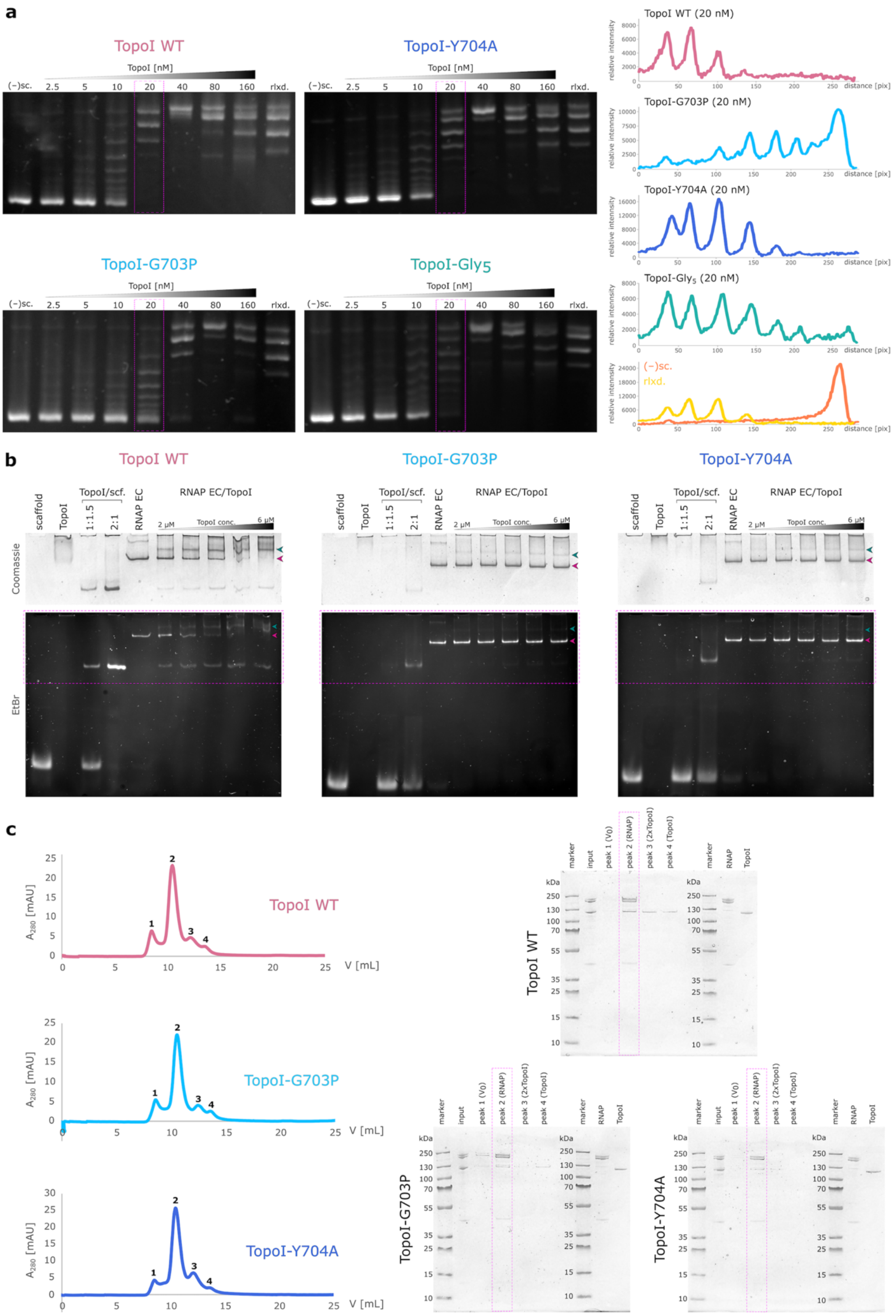

**Supplementary figure 6: *In vitro* assays on TopoI mutants. a**, DNA relaxation assay. Supercoiled pUC19 plasmid was relaxed in presence of increasing concentrations of TopoI (wildtype or mutant). The products were separated on a 1 % agarose gel in absence of intercalators and subsequently stained with ethidium bromide. The lanes corresponding to 20 nM TopoI were plotted as histograms (right). (–)sc. – negatively supercoiled plasmid, rlxd. – relaxed topoisomers. **b**, Electromobility shift assay of RNAP-TopoI complexes with TopoI mutants. RNAP EC was assembled using duplex scaffold and incubated with increasing amounts of TopoI (wt or mutants). Complexes corresponding to RNAP EC and RNAP EC-TopoI are highlighted with pink and green arrows in gels stained with Coomassie (top) or ethidium bromide (bottom), respectively. Both tested TopoI mutants (G703P and Y704A) show weaker binding to RNAP EC compared to wildtype. **c**, Size exclusion chromatography of RNAP-TopoI complexes with TopoI mutants (G703P and Y704A) on duplex scaffolds. In all cases TopoI co-elutes with RNAP (peak 2), but the amount of mutant TopoI is lower compared to the wt (top vs. bottom).

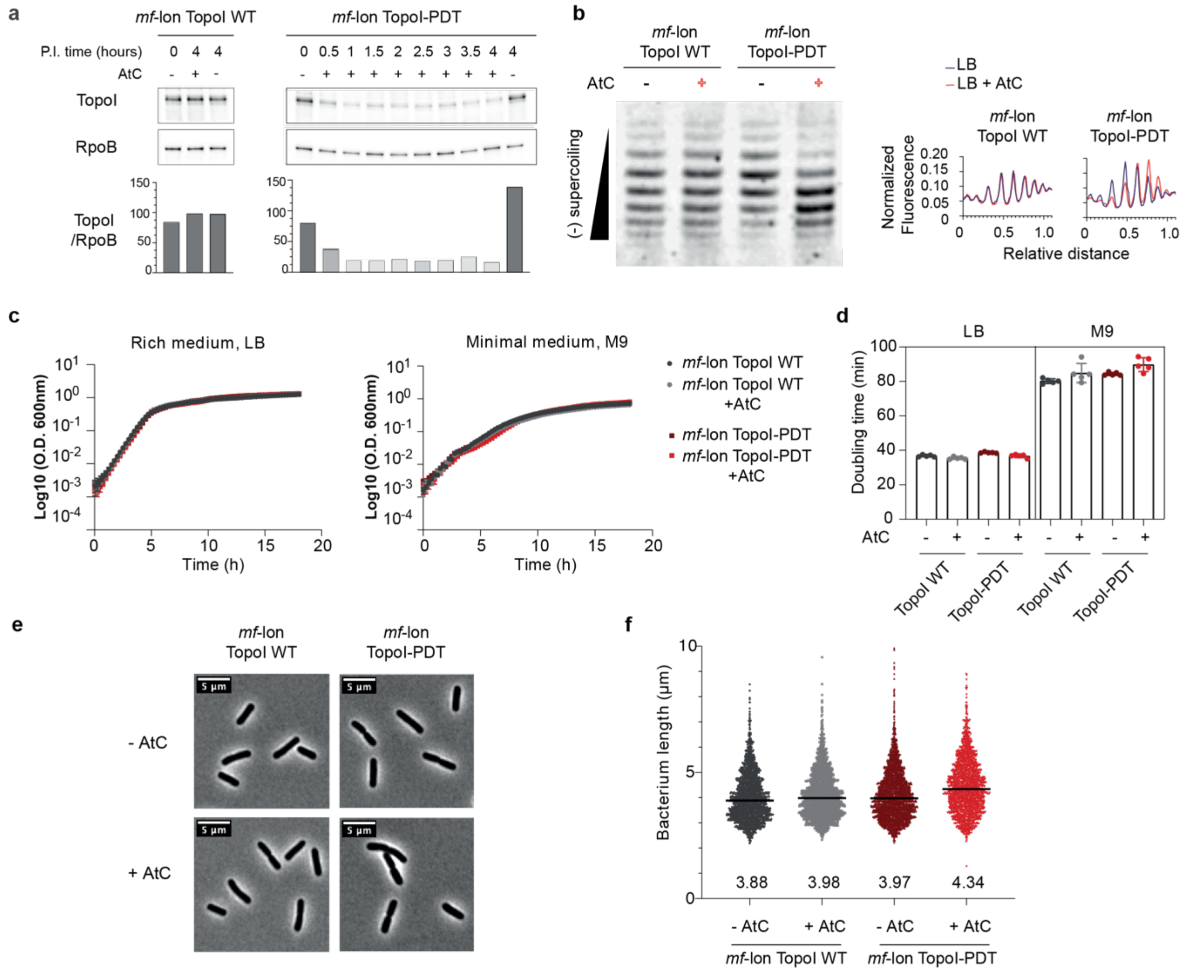

**Supplementary figure 7: Study of TopoI-PDT depletion.** **a**, Measurement of TopoI levels by western blot in *mf*-lon TopoI WT and *mf*-lon TopoI-PDT cells with anti-TopoI antibody. Western-blot quantification showed a degradation of TopoI levels up to 85% one hour post-induction (P.I.). **b**, One-dimensional chloroquine-agarose gel electrophoresis of pUC18 isolated from *mf*-lon TopoI WT or *mf*-lon TopoI-PDT. In this condition, supercoiling increases migration speed. The fluorescence intensity was measured and compared between the two conditions. **c**, Growth of MG1655 *mf*-lon TopoI WT and TopoI-PDT strains in LB and M9 minimal medium in absence or in presence of AtC. $OD_{600}$ was measured in Tecan Spark Microplate Reader within a 96 well plate. **d**, Corresponding doubling times were calculated for each well. The mean and standard deviation obtained are represented. **e**, Microscopy imaging of the strains grown in LB medium and fixed after four hours post-induction or not with AtC (50 ng/mL). **f**, Bacteria length was measured with a custom macro in the Fiji software.

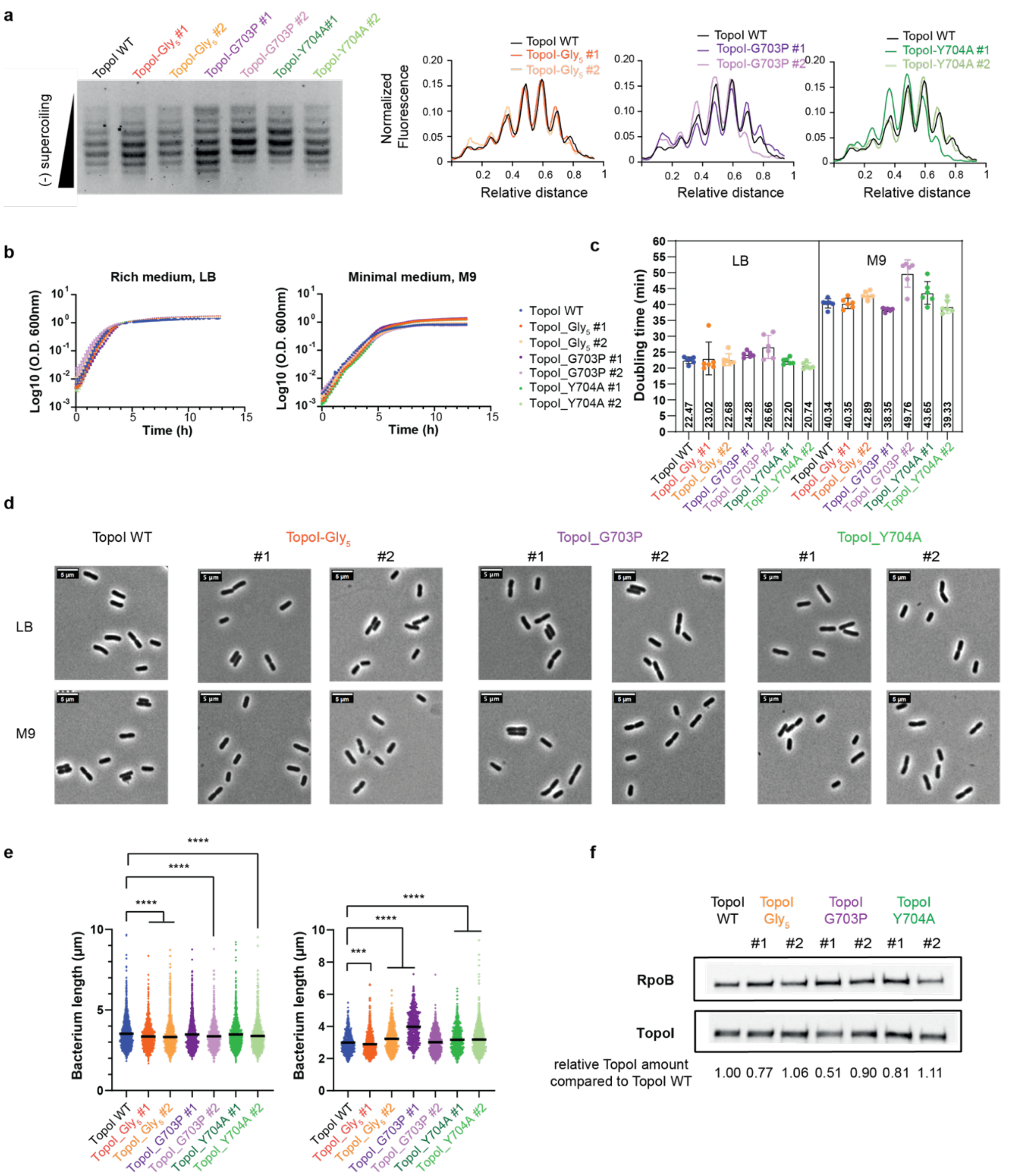

**Supplementary figure 8: Study of CRISPR-Cas9 mediated point mutations of Topo I. a**, One-dimensional chloroquine-agarose gel electrophoresis of pUC18 isolated from TopoI WT or mutants. At the chloroquine concentration employed, migration speed increases with supercoiling levels. Band intensity was evaluated using ImageJ and is shown on the right. **b**, Growth of *E. coli* MG1655 strains harbouring WT or mutant versions of TopoI in LB and M9 minimal medium. $OD_{600}$ was measured in Tecan Spark Microplate Reader within a 96 well plate. **c**, Corresponding doubling times were calculated for each well. The mean and standard deviation obtained are represented. **d**, Microscopy imaging of the strains in LB and M9 minimal medium. The cells were fixed during the exponential growth phase ($OD_{600}$≈0.3). **e**, Bacteria length was measured with a custom macro in the Fiji software. The significance of the two-tailed Mann-Whitney test between average of conditions is indicated by stars (*: <0.032; **: <0.0021; ***:<0.0002; ****: <0.0001). **f**, Measurement of TopoI amount by western blot in *E. coli* MG1655 strains harbouring WT or mutant versions of TopoI with an anti-TopoI antibody.

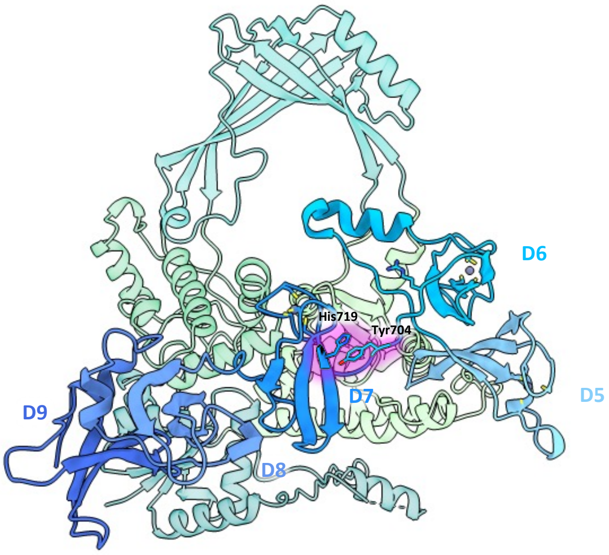

**Supplementary movie 1: TopoI-CTD rotation.** The TopoI-CTD exhibits a complex rotational movement relative to the catalytic core in comparison to the crystal structure (PDB ID 4RUL). It rotates around residue Gly703 in the linker connecting domains D6 and D7, and it rotates around residue Lys752 in the linker connecting domains D7 and D8.

>> https://drive.google.com/file/d/1JqhGX6ss5xGvqEA2eNxt9ScdVgo6U02L/view?usp=drive_link

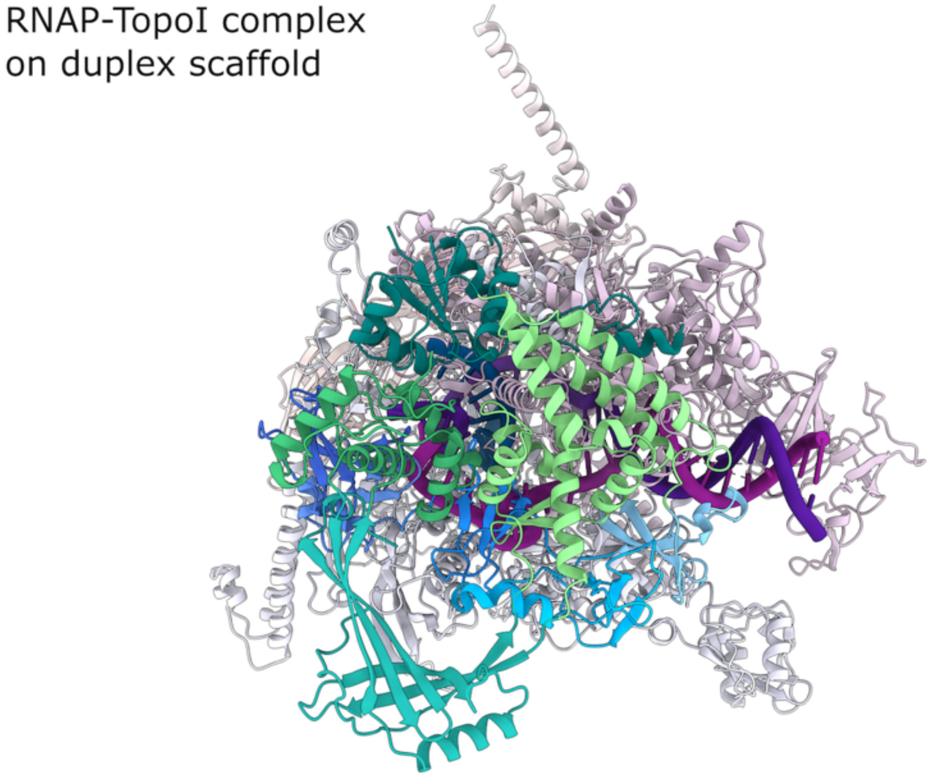

**Supplementary movie 2: RNAP-TopoI complex orientations.** This movie is a morph to illustrate and compare the different relative orientations that TopoI adopts relative to RNAP in the complex on the duplex scaffold (relaxed DNA), bubble scaffold (consensus reconstruction), and long-overhang scaffold (orientation 1 followed by orientation 2).

>> https://drive.google.com/file/d/1_-hlgKVa2g0deoW1i7uIgCFYEPntly_-/view?usp=drive_link

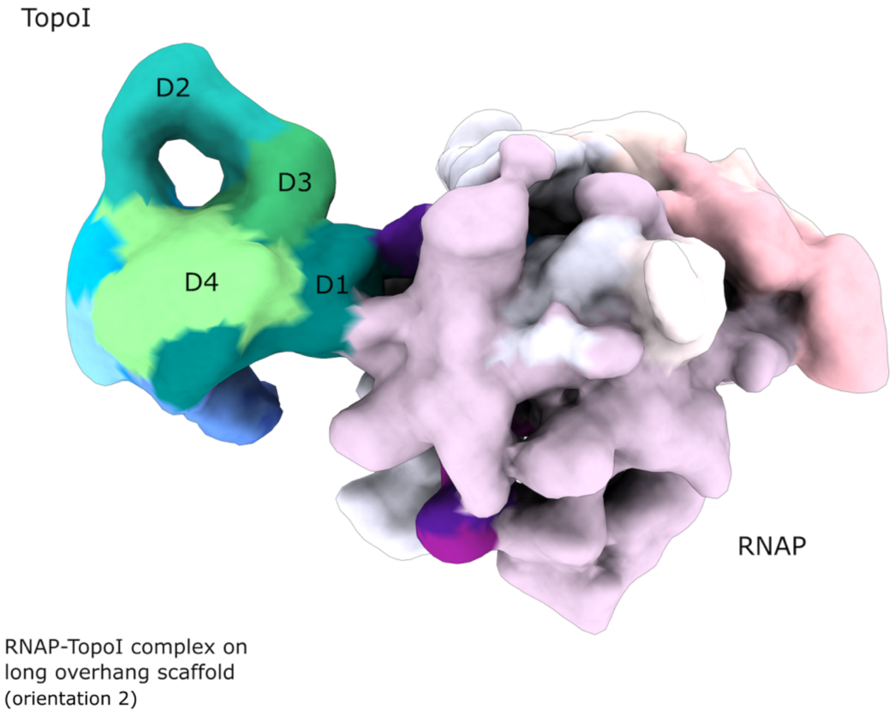

**Supplementary movie 3: 3D variability analysis of TopoI gate opening on the long overhang scaffold.** This movie shows a series of reconstructions as obtained by 3D variability analysis in CryoSPARC to illustrate the conformational variability of TopoI amongst the particles corresponding to orientation 2 in the RNAP-TopoI complex on the long-overhang scaffold.
>> https://drive.google.com/file/d/1umxpZjB5ddfdkFklnhOWYEPFOVw00Bxv/view?usp=drive_link

**Supplementary table 1:** Data collection and refinement statistics for RNAP-TopoI complexes.

| **Data collection** | **RNAP-TopoI complex on duplex scaffold** | **RNAP-TopoI complex on bubble scaffold – consensus reconstruction** | **RNAP-TopoI complex on bubble scaffold – "two-bubble" configuration** | **RNAP-TopoI complex on bubble scaffold – "single-bubble" configuration** | **RNAP-TopoI complex on long overhang scaffold – orientation 1**** | **RNAP-TopoI complex on long overhang scaffold – orientation 2**** |
|---|---|---|---|---|---|---|
| Particles | 113760 | 49760 | 35119 | 14641 | 13653 | 10189 |
| Pixel size (Å) | 0.839 | 0.862 | 0.862 | 0.862 | 0.645 | 0.645 |
| Defocus range (µm) | -0.8 – -2.5 | -0.8 – -2.5 | -0.8 – -2.5 | -0.8 – -2.5 | -0.8 – -2.0 | -0.8 – -2.0 |
| Voltage (kV) | 300 | 300 | 300 | 300 | 300 | 300 |
| Total electron dose ($e^- Å^{-2}$) | 56.60 | 54.03 | 54.03 | 54.03 | 49.91 | 49.91 |
| Collected at | Titan Krios with K3 detector (ESRF Grenoble, France, February 2022)[34] | Titan Krios with K3 detector (FRISBI / Instruct-ERIC platform, IGBMC-CBI Illkirch, France, July 2021) | | | Titan Krios with K3 detector (EMBL Heidelberg, Germany, April 2021) | |
| **Model composition** | | | | | | |
| Non-hydrogen atoms | 33857 | 33365 | 33365 | 33385 | 33652 | 33510 |
| Protein residues | 4084 | 4084 | 4084 | 4084 | 4084 | 4065 |
| Nucleotide residues | 89 | 65 | 65 | 66 | 79 | 79 |
| Ligands ($Zn^{2+}$/$Mg^{2+}$) | 5/1 | 5/1 | 5/1 | 5/1 | 5/1 | 5/1 |
| **Model vs. data** | | | | | | |
| Nominal resolution (Å) | 2.7 | 3.8 | 4.3 | 4.3 | 4.0 | 3.9 |
| Map-sharpening B-factor ($Å^2$) | N/A* | 103.5 | / | / | 84.6 | 70.6 |
| Map cross-correlation (within mask) | 0.83 | 0.85 | 0.81 | 0.79 | 0.83 | 0.84 |
| Average B factor protein ($Å^2$) | 71.28 | 368.47 | 390.37 | 332.95 | 274.73 | 260.49 |
| Average B factor nucleotide ($Å^2$) | 109.18 | 242.00 | 275.18 | 280.66 | 292.73 | 246.54 |
| **RMS deviations** | | | | | | |
| Bond lengths (Å) | 0.010 | 0.006 | 0.006 | 0.007 | 0.010 | 0.006 |
| Bond angles (°) | 0.642 | 1.094 | 1.099 | 0.851 | 1.121 | 0.865 |
| **Ramachandran plot** | | | | | | |
| Favoured (%) | 98.55 | 97.00 | 97.00 | 96.67 | 96.73 | 96.17 |
| Allowed (%) | 1.45 | 2.95 | 2.88 | 3.13 | 3.27 | 3.71 |
| Outliers (%) | 0.00 | 0.05 | 0.12 | 0.20 | 0.00 | 0.12 |
| **Validation** | | | | | | |
| Molprobity Score | 1.32 | 1.81 | 1.85 | 2.13 | 1.89 | 2.38 |
| Molprobity Clash score | 5.83 | 12.25 | 13.04 | 15.83 | 15.10 | 20.53 |
| Rotamer outliers (%) | 0.03 | 1.12 | 1.15 | 1.90 | 0.52 | 2.58 |
| **Data availability** | | | | | | |
| PDB accession number | 9GDA | 9GDB | 9GDD | 9GDC | 9GDE | 9GDH |

| | | | | | | |
|---|---|---|---|---|---|---|
| EMDB accession number | EMD-51259<br>EMD-51252<br>EMD-51253<br>EMD-51254<br>EMD-51255<br>EMD-51256<br>EMD-51257<br>EMD-51258<br>EMD-51267<br>EMD-51265<br>EMD-51266 | EMD-51260 | EMD-51262 | EMD-51261 | EMD-51263 | EMD-51264 |

* Map was sharpened using DeepEMhancer [35].

** RNAP-TopoI complex on long overhang scaffold in orientation 3 and with an open TopoI-gate were deposited in the EMDB (IDs: EMD-51268, and EMD-51269, respectively)

### Supplementary table 2: Protein-protein contacts between RNAP and TopoI.

| RNAP-TopoI complex | local resolution for TopoI at FSC = 0.5 (Å) | RNAP | TopoI |
|---|---|---|---|
| complex on duplex scaffold | 3.3 | β protrusion (residues 31 – 139 and 456 – 512) | domain D7 (residues 707 – 750) |
| | | β′ tip of the clamp domain (residues 127 – 196) | domain D5 (residues 591 – 635) and TOPRIM insertion loop (residues 37 – 60)* |
| complex on short overhang scaffold – orientation 1 | 5.0 | β′ coiled coil (residues 264 – 332) | TOPRIM domain D1 (residues 1 – 157) |
| | | β′ Zn finger (residues 63 – 95) | CAP domain D3 (residues 279 – 404) |
| complex on short overhang scaffold – orientation 2 | 7.3 | β tip of the SI2 domain (residues 943 – 1037) | CAP domain D3 (residues 279 – 404) |
| | | β protrusion (residues 31 – 139, 456 – 512) | domain D9 (residues 825 – 865) |
| complex on bubble scaffold | 6.4 | β′ tip of the clamp domain (residues 127 – 196) | loop in domain D2 (442 – 448) |
| | | β′ coiled coil (residues 264 – 332) | domain D7 (residues 707 – 750) |
| | | β flap (residues 829 – 937 and 1040 – 1059) and β′ Zn finger (residues 63 – 95) | domain D9 (residues 825 – 865) |
| complex on long overhang scaffold – orientation 1 | 4.8 | β lobe (residues 154 – 447) | CAP domain D3 (residues 279 – 404) |
| | | β′ tip of the clamp domain (residues 127 – 196) | CAP domain D3 (residues 279 – 404) and loop in domain D2 (442 – 448) |
| complex on long overhang scaffold – orientation 2 | 4.5 | β′ coiled coil (residues 264 – 332) | TOPRIM domain D1 (residues 1 – 157) |
| complex on long overhang scaffold – orientation 3 | 7.8 | β protrusion (residues 31 – 139 and 456 – 512) | TOPRIM domain D1 (residues 1 – 157) |
| | | β lobe (residues 154 – 447) and β′ tip of the clamp domain (residues 127 – 196) | CAP domain D3 (residues 279 – 404) |

* The contact was observed only in a subset of particles.

### Methods

#### *E. coli* RNAP purification

Two plasmids, pVS11_rpoA_rpoB_rpoC_HRV3C_His10_rpoZ (encoding all subunits of *E. coli* core RNAP with a $His_{10}$-tag on the CTD of β' subunit) and pACYC_Duet1_rpoZ (to avoid its sub-stoichiometric amounts of ω subunit), were co-transformed into LACRII expression strain (a derivative of *E. coli* LOBSTR[36]). The proteins were expressed by inducing a 6 L culture (100 µg/mL ampicillin and 34 µg/mL chloramphenicol) grown at 37°C at $OD_{600}$=0.6-0.8 with 1 mM IPTG. After 2 hours cells were harvested by centrifugation at 4500 g, 4 °C for 30 min. The pellet was resuspended in 5 volumes of lysis buffer (50 mM Tris/HCl pH 8.0, 5 % glycerol, 1 mM EDTA pH 8.0, 10 mM DTT, 0.1 mM PMSF, 1 mM benzamidine, 10 µM $ZnCl_2$) supplemented with cOmplete EDTA-free protease inhibitor cocktail (Sigma Aldrich, 1 tablet/50 mL) and DNAse I (~0.1 mg/50 g cell pellet), and lysed by sonication. The lysate was centrifuged at 40000 g, 4 °C for 30 min and RNAP was isolated from the soluble fraction by polyethyleneimine (PEI) fractionation followed by ammonium sulfate precipitation as described previously[37]. The precipitate was recovered by centrifugation at 40 000g, 4 °C for 30 min, dissolved in 50 – 60 mL of IMAC buffer A (20 mM Tris/HCl pH 8.0, 1 M NaCl, 5 % glycerol, 5 mM β-mercaptoethanol, 0.1 mM PMSF, 1 mM benzamidine, 10 mM $ZnCl_2$) and loaded on a pre-equilibrated 20 mL Ni-IMAC Sepharose HP column (GE Healthcare) using an ÄKTA system. The column was washed with 2 CV of buffer A, then 2 CV of 2 % IMAC buffer B (20 mM Tris/HCl pH 8.0, 1 M NaCl, 5 % glycerol, 5 mM β-mercaptoethanol, 0.1 mM PMSF, 1 mM benzamidine, 10 mM $ZnCl_2$, 250 mM imidazole) followed by a gradient to 16 % B over 1 CV, and finally 5 CV of 16 % B. The RNAP was eluted in 3 CV of 100 % B, and the peak fractions were pooled. The His-tag was cleaved with HRV3C protease (1 mg per 8 mg of RNAP) during an overnight dialysis against the dialysis buffer (20 mM Tris/HCl pH 8.0, 1 M NaCl, 5 % glycerol, 5 mM β-mercaptoethanol, 10 µM $ZnCl_2$). The sample was reloaded on the Ni-IMAC Sepharose HP column to separate the cleaved RNAP from the non-cleaved RNAP and HRV3C protease. The cleaved RNAP was then dialysed into BioRex buffer C (10 mM Tris/HCl pH 8.0, 5% glycerol, 0.1 mM EDTA, 1 mM DTT, 0.1 mM PMSF, 1 mM benzamidine, 10 µM $ZnCl_2$) twice to decrease the conductivity below 10 mS/cm and loaded onto a pre-equilibrated BioRex 70 column (Biorad). The column was washed with 2 CV of BioRex buffer C and RNAP was eluted using a gradient 0 – 100 % BioRex buffer D (10 mM Tris/HCl pH 8.0, 5% glycerol, 0.1 mM EDTA, 1 mM DTT, 0.1 mM PMSF, 1 mM benzamidine, 10 µM $ZnCl_2$, 1 M NaCl) over 5 CV. The peak fractions were pooled, concentrated and then purified by size exclusion chromatography on a HiLoad Superdex 200 PG 26/600 column (GE Healthcare), pre-equilibrated with GF buffer (10 mM HEPES pH 8.0, 0.5 M KCl, 1 % glycerol, 2 mM DTT, 0.1 mM PMSF, 1 mM benzamidine, 10 µM $ZnCl_2$, 1 mM $MgCl_2$). Peak fractions were pooled and dialysed into the EM buffer (10 mM HEPES/KOH pH 7.5, 150 mM KOAc, 5 mM $Mg(OAc)_2$, 10 µM $ZnCl_2$, 2 mM DTT). RNAP was then concentrated, aliquoted and flash-frozen in liquid nitrogen. The aliquots were stored at -80 °C until use.

#### *E. coli* TopoI purification

A plasmid encoding either the wild type *E. coli* TopoI or the mutant protein (pAX7-EcTopI) with the N-terminal cleavable $His_{10}$-tag followed by the TwinStrep tag was transformed into LACRII expression strain. The protein was expressed by inducing a 3 L culture (50 µg/mL kanamycin) grown at 37°C at $OD_{600}$=0.7 with 0.8 mM IPTG for 3 h, and then harvested by centrifugation at 4000 g, 4°C for 20 min. The pellets were resuspended in 5 volumes of cold lysis buffer (20 mM Tris/HCl pH 7.0 at RT, 500 mM NaCl, 5 % glycerol, 10 mM imidazole, 10 µM $ZnCl_2$, 0.1 mM PMSF, 1 mM benzamidine, 6 mM β-mercaptoethanol) with added cOmplete Protease inhibitor cocktail (Roche; 1 tablet per 50 mL buffer) and DNase (~20 µg per 1 g of pellet) and lysed by sonication. The bacterial lysate was centrifuged at 45000 g, 4°C for 45 min, and filtered through 0.22 µm filter. The lysate was loaded on two 5 mL HiTrap IMAC HP columns pre-equilibrated with the lysis buffer using the ÄKTA system. The columns were washed with 10 CV lysis buffer, followed by 10 CV 5 % IMAC elution buffer (20 mM Tris/HCl pH 7.0 at RT, 500 mM NaCl, 5 % glycerol, 500 mM imidazole, 10 µM $ZnCl_2$, 0.1 mM PMSF, 1 mM benzamidine, 6 mM β-mercaptoethanol) and then eluted with 15 CV 100 % IMAC elution buffer. The $His_{10}$-tag was cleaved with His-tagged HRV3C protease (~ 1 mg per 5 mg of TopoI) during a one-hour dialysis at 4 °C into dialysis buffer (20 mM Tris/HCl pH 7.0 at RT, 500 mM NaCl, 5 % glycerol, 10 mM imidazole, 10 µM $ZnCl_2$, 6 mM β-mercaptoethanol). Afterwards, the sample was re-loaded on the HiTrap IMAC HP column to separate the cleaved TopoI from the uncleaved and the HRV3C protease. The column was washed with 10 CV lysis buffer. The flowthrough containing cleaved TopoI was diluted with dilution buffer (20 mM Tris/HCl pH 7.0 at RT, 5 % glycerol, 10 µM $ZnCl_2$, 0.1 mM EDTA, 0.1 mM PMSF, 1 mM benzamidine, 1 mM DTT) to a final salt concentration 150 mM NaCl and loaded on a 5 mL HiTrap Heparin HP column that had been pre-equilibrated with Hep buffer (20 mM Tris/HCl pH 7.0 at RT, 150 mM NaCl, 5 % glycerol, 10 µM $ZnCl_2$, 0.1 mM EDTA, 0.1 mM PMSF, 1 mM benzamidine, 1 mM DTT). The column was washed with 5 CV Hep buffer followed by a gradient elution 0 – 100% Hep elution buffer (20 mM Tris/HCl pH 7.0 at RT, 1 M NaCl, 5 % glycerol, 10 µM $ZnCl_2$, 0.1 mM EDTA, 0.1 mM PMSF, 1 mM benzamidine, 1 mM) over 20 CV. For structural studies, TopoI was additionally purified by size exclusion chromatography on a HiLoad 16/600 Superdex 200 pg column (GE Healthcare), pre-equilibrated with GF buffer (10 mM HEPES/KOH pH 7.5, 500 mM KOAc, 5 mM $Mg(OAc)_2$, 5 % glycerol, 10 µM $ZnCl_2$, 1 mM DTT). The protein was concentrated, aliquoted and flash-frozen in liquid nitrogen, then stored at -80°C until use.

**Reconstitution of nucleic acid scaffolds**

A nucleic acid scaffold was assembled from short DNA and RNA oligonucleotides that mimic a transcription bubble. The central part of tDNA and ntDNA contain mismatches to allow annealing of the RNA (Fig. 2a).
DNA (TriLink, Sigma Aldrich, or Eurofins) and RNA (Dharmacon) oligonucleotides were chemically synthetised and purified by the manufacturer. RNA was deprotected following the protocols provided by the manufacturer. Both DNA and RNA were dissolved in RNase free water and stored at -80°C. For a 100 µM scaffold stock, the respective RNA, tDNA and ntDNA oligonucleotides were mixed in equimolar concentration (100 µM each) in RB buffer (10 mM Tris/HCl pH 7.0, 40 mM NaCl, 5 mM $MgCl_2$), and then annealed in a thermocycler using the following protocol: 95°C for 2 min, 75°C for 2 min, 45°C for 5 min, then -1°C/min until 4°C. The reconstituted scaffolds were stored at -20 °C until use.

**RNAP-TopoI complex analysis by size exclusion chromatography**

To prepare RNAP EC, purified RNAP (final concentration 20 µM) and a nucleic acid scaffold (final concentration 24 µM) were mixed in EM buffer (10 mM HEPES/KOH pH 7.5, 150 mM KOAc, 5 mM $Mg(OAc)_2$, 10 µM $ZnCl_2$, 2 mM DTT) incubated at 37 °C for 10 min and purified on a Superdex 200 Increase 10/300 GL (GE Healthcare) column equilibrated with EM buffer. To prepare RNAP-TopoI complex, RNAP EC (final concentration 5 µM) and TopoI (final concentration 6 µM) were mixed in EM buffer and incubated at 37°C for 5 min. The sample was separated on a Superdex 200 Increase 10/300 GL column as described and peak fractions were analysed by SDS-PAGE.

**RNAP-TopoI complex analysis by gel shift assay**

RNAP EC was prepared and purified by size-exclusion chromatography as described before. To prepare the samples for the gel shift assay, RNAP EC (2 µM final concentration) was mixed with TopoI (final concentration 2 – 6 µM) in EM buffer (10 mM HEPES/KOH pH 7.5, 150 mM KOAc, 5 mM $Mg(OAc)_2$, 10 µM $ZnCl_2$, 2 mM DTT) and incubated at 37 °C for 5 min. The samples were mixed with 2x native loading dye (250 mM Tris/HCl pH 8.8 at 4 °C, 0.88 M L-alanine, 10 % glycerol, 0.25 % (w/v) bromophenol blue, 0.25 % (w/v) xylene cyanol) and analysed on a 6 % native polyacrylamide gel at 4 °C. After the pre-run (10 mA, 30 min), the samples were separated at 120 V, 65 – 75 min in Tris-alanine running buffer (250 mM Tris/HCl pH 8.8 at 4 °C, 0.88 M L-alanine). Each gel was run in a duplicate – the first was stained with ethidium bromide and the second gel with Coomassie.

**Sample preparation for cryo-EM**

RNAP EC was prepared in EM buffer (10 mM HEPES/KOH pH 7.5, 150 mM KOAc, 5 mM $Mg(OAc)_2$, 10 µM $ZnCl_2$, 2 mM DTT) in a final volume of 150 µL by incubating RNAP (final concentration 50 µM) and a nucleic acid scaffold (final concentration 75 µM) at 37 °C for 10 min, and then purified by size exclusion chromatography on a Superose 6 10/300 GL column (Cytiva) in EM buffer. To prepare RNAP-TopoI complex, the purified RNAP EC and TopoI were mixed in an equimolar ratio in EM buffer and incubated at 37 °C for 5 min. Prepared RNAP-TopoI sample was centrifuged at 20000g, 4 °C for 10 min to remove aggregates and transferred to a fresh tube. The concentration was estimated on a Nanodrop using 1 $A_{280}$ = 1 mg/mL protein mode. When necessary, the concentration was adjusted to approximately 10 mg/mL with EM buffer, and CHAPSO was added to 8 mM final concentration. The quality of each sample was verified on a 4 – 15 % gradient PhastGel under native conditions, using a commercial Phast System (Cytiva). UltrAuFoil Holey Gold R1.2/1.3 Au 300 mesh grids were glow-discharged using a plasma cleaner (Fischione 1070) with a gas mixture of $Ar/O_2$ (90/10) at 70 % power, for 35 s. Freezing was done using a Vitrobot Mark IV at 10 °C and 95 – 100 % chamber humidity. 3 µL of sample was applied on a grid, which was then blotted for 2 s with force 8 before being plunged into liquid ethane. Frozen grids were stored in liquid nitrogen.

**Cryo-EM data collection and processing**

The datasets were collected on Titan Krios transmission electron microscopes controlled by SerialEM (ver. 3.6)[38] as described in Supplementary table 1. Where on-the-fly processing was available (Titan Krios, CBI-IGBMC), motion correction, CTF estimation and particle picking were done in Warp[39]. At ESRF, only motion correction was performed on-the-fly using MotionCor2[40]. Otherwise, motion correction and CTF estimation were done in cryoSPARC[21] using their own implementation, and particles were picked in crYOLO with the pre-trained general model[41]. Data was further processed in cryoSPARC[21]. The initial clean-up after particle extraction was done by 2D classification, and a subset of the best-looking 2D classes was used to obtain an *ab-initio* model. Multiple rounds of heterogeneous refinement were performed to further remove low-quality particles and to obtain starting models for different conformations of RNAP-TopoI complexes. To better separate various conformations and find more homogeneous subsets of particles that refine to higher resolution, several rounds of 3D variability analysis[42] were performed with a wide, soft mask around TopoI, and using filter resolution of 10 – 15 Å. Results were visualised using either clustering mode or intermediate frames, both of which make 3D reconstructions from particle images using the input alignments. Particles giving rise to poor reconstructions were discarded, whereas those from similar-looking reconstructions were fed into subsequent rounds of refinement and 3DVA. All final maps were refined using non-uniform refinement[43]. For the RNAP-TopoI complex on duplex scaffold, a local refinement was performed (using alignment prior and with non-uniform refinement option enabled) on several regions: whole TopoI, D2/D3 of TopoI, RNAP core, RNAP swivel module, and β jaw of RNAP. The resulting maps were sharpened with DeepEMhancer[35] and combined into one map using Phenix[44]. Initial placement of pdb models (6ALH for RNAP[19], 4RUL for TopoI[9]) into the EM maps was done manually in UCSF Chimera[45], and nucleic acid scaffolds were built in Coot[46]. The RNAP-TopoI complex on duplex scaffold was refined against the EM map in Phenix using secondary structure, Ramachandran and base pair restraints. Coot and Isolde[47] were then used to manually rebuild regions of the model that were in disagreement with the EM map, and to optimize the geometry of the model (remove Ramachandran outliers, optimize rotamers, fix steric clashes). The rebuilt model was then again refined in Phenix. For the RNAP-TopoI complex on the bubble scaffold, RNAP was refined in Phenix (with the secondary structure, Ramachandran and base pair restraints), and rigid body fitting of TopoI NTD and CTD into the lowpass-filtered maps were performed in Chimera. For the complex on long overhang scaffold, RNAP was refined in Phenix (with the secondary structure, Ramachandran and base pair restraints), and flexible fitting of TopoI (using distance restraints for each domain) was performed with Isolde[47] against lowpass-filtered maps. The rotation angles for TopoI-CTD were measured in Pymol[48] using a script for drawing rotation axis (https://pymolwiki.org/index.php/RotationAxis). The figures were prepared with UCSF Chimera[45] and ChimeraX[49].

### DNA cleavage assay on RNAP-TopoI complex

Nucleic acid scaffolds were reconstituted using tDNA with a phosphorylated 5′-end. RNAP EC complex was prepared in EM buffer (10 mM HEPES/KOH pH 7.5, 150 mM KOAc, 5 mM $Mg(OAc)_2$, 10 µM $ZnCl_2$, 2 mM DTT) and purified by size exclusion chromatography on a Superose 6 10/300 GL column (Cytiva) using EM buffer lacking $Mg^{2+}$ as described above. Fractions containing RNAP EC were pooled and concentrated. The ntDNA strand in RNAP EC was radioactively labelled on the 5′-end with $^{32}$P-γ-ATP (10 mCi/mL). 10 µM purified RNAP EC, 2.5 µL $^{32}$P-γ-ATP and 0.5 U/µL T4 PNK (Invitrogen) were mixed in the PNK buffer A (50 mM Tris/HCl pH 7.6, 10 mM $MgCl_2$, 150 mM KOAc, 5 mM DTT, 0.1 mM spermidine) in a final volume of 10 µL and incubated at 37 °C for 45 min. Cold ATP was subsequently added into the reaction mixture to a final concentration of 2.5 µM, further incubated for 15 min, then stopped by addition of 25 mM EDTA. Labelled RNAP EC was then used to spike a 10x stock for DNA cleavage assay. 0.5 µM was mixed with 9.5 µM of non-labelled RNAP EC in the EM buffer without $Mg^{2+}$ to give total final concentration 10 µM. For a control reaction, ntDNA oligonucleotide was labelled with $^{32}$P-γ-ATP as described for RNAP EC. Labelling reaction was

terminated by incubation at 95 °C for 5 min. Excess ATP was removed by passing the reaction mixture through a BioRad column filled with 10 volumes of Sephadex G-50 resin. Labelled ntDNA was then used to reconstitute the respective nucleic acid scaffold. For 10 µM stock, 12 µM RNA, 12 µM tDNA, 9.5 µM non-labelled ntDNA and 0.5 µM labelled ntDNA were mixed in RB buffer (10 mM Tris/HCl pH 7.0, 40 mM NaCl, 5 mM $MgCl_2$). The mixture was heated to 95 °C for 2 min and then immediately placed into a beaker filled with hot water (> 80 °C), where it was left to cool to room temperature. A set of synthetic single-stranded DNA oligonucleotides (5 nt, 10 nt, 15 nt, and 20 nt) were radioactively labeled and used a marker. For DNA cleavage assay, TopoI was diluted with EM buffer without $Mg^{2+}$ to 10 µM working concentration. 2 µL of TopoI (final concentration 1 µM) was added to 18 µL of spiked RNAP EC (or spiked ntDNA or labelled scaffold for control reactions; final concentration 1 µM) in DNA cleavage buffer (10 mM HEPES/KOH pH 7.5 at RT, 150 mM KOAc, 5 mM EDTA, 10 µM $ZnCl_2$, 2 mM DTT). Reaction mixtures were incubated at 37 °C for 1 h. After this time, 10 µL aliquot was removed and mixed with equal volume of stop solution (80 % formamide, 0.2 M NaOH, 0.04 % bromophenol blue). Then, 2.5 µL of 5x Mg/NaCl solution (50 mM $MgCl_2$, 5 M NaCl) was added to the remaining volume and incubated at room temperature for 5 min. Again, 10 µL aliquot was taken and mixed with stop solution. All samples were boiled at 95 °C for 5 min before being loaded on the gel (1 – 5 µL/lane).
Samples were analysed on 15 % denaturing PAGE gel. The gel was pre-run for 20 – 30 min at 50 W in 1x TBE buffer and then run for 1.5 h after loading the samples. Afterwards, the gel was transferred into an exposure cassette and exposed at -80 °C overnight. The phosphor screen was scanned using a Typhoon phosphorimager (GE Healthcare).

**Cloning of TopoI mutants**

The plasmids for expression of TopoI mutants (G703P, Y704A and $Gly_5$) were generated from the pAX7-EcTopI plasmid by *In Vivo* Assembly cloning[50]. The entire plasmid was amplified by inverse PCR with the mutagenic primers and Phusion DNA-polymerase (Invitrogen) by the manufacturer's instructions. Afterwards, the PCR template was digested with 20 U of DpnI (NEB) at 37 °C for 1 h, and 10 µL of the reaction mixture were transformed into *E. coli* TOP10 competent cells (Invitrogen). Plasmids were isolated from the single colonies and the presence of the desired mutations was confirmed by DNA sequencing.

**DNA relaxation assay**

Supercoiled pUC19 plasmid was purified from an overnight culture of *E. coli* TOP10 strain with a NucleoBond Xtra Maxi kit (Macherey-Nagel) according to the manufacturer's instructions. Supercoiled pUC19 plasmid was diluted in DNA relaxation buffer (20 mM Tris/acetate pH 7.9 at RT, 100 mM KOAc, 10 mM $Mg(OAc)_2$, 10 µM $ZnCl_2$, 2 mM DTT, 0.1 mg/mL acetylated BSA) to concentration 11.1 ng/µL. The reaction was set up on ice, and the components were added to the tube in the following order: 2 µL TopoI (final concentrations 2.5 – 160 nM) and 18 µL pUC19 (200 ng of plasmid/reaction; final concentration 6 nM). The samples were incubated at 37 °C for 30 min and the reaction terminated by addition of 1 % SDS. The samples were analysed on 1 % agarose gel run in 1x TBE buffer at 90 V for 3 h. The gel was stained with ethidium bromide (2 µg/mL in 1x TBE) for 15 min and de-stained in water for 5 – 10 min then imaged under the UV light. Band intensities were evaluated using Fiji[51].

**Bacteria, plasmids and growth conditions**

Strains used in this study are derived from MG1655 *E. coli* strain. TopoI depletion was performed in the *mf*-lon strain described in[24] with topA gene fused with the sequence encoding the PDT3A tag. Mutations $Gly_5$, G703P and Y704A were inserted in chromosome using Cas/pTargetF system described in[52].

The low-copy plasmid, pGB2-Para, used for complementation assay of TopoI-PDT strain was designed to overexpress TopoI-WT, TopoI-$Gly_5$, TopoI-G703P or TopoI-Y704A under the control of the arabinose promoter. Plasmid pMLS-AAV used for expression assay was derived from a genetic construct with pSC101 origin of replication[26]. In this study, mCerulean and mVenus were tagged in C-terminal with AAV peptide to reduce their half-life down to 1 hour. pUC18 was used for plasmid supercoiling measurements.

All strains were grown in M9 minimal medium using the following recipe: 1X M9 Minimal Salts (from Sigma Aldrich); 0.4 % D-glucose; 1 % Casaminoacids; 2 mM $MgSO_4$ and 0.1 mM $CaCl_2$ or in LB rich medium. When needed, inducers were used at 0.2 % for L-arabinose and 50 ng/mL for AtC (anhydrotetracycline).

**Antibodies**

Rabbit polyclonal antibodies were produced by Agro-Bio using full length recombinant EcTopoI as an immunogen and the serum was used at 1/1000 dilution. Secondary antibody was purchased from Jackson ImmunoResearch (#711-035-152) and used at 1/10000 dilution. Anti RNApol Beta antibody was purchased from Biolegend (#662904) and used at the 1/5000 dilution.

**Construction of pMLS-AAV**

The pMLS plasmid was amplified by PCR with two pairs of primers (mCer-Ter-fwd/mVen-End-rev and mVen-Ter-fwd/mCer_End_rev), to obtain two fragments covering the full length plasmid. The AAV peptide was amplified on an existing matrix using pairs of primers with flanking sequences corresponding to pMLS: mCer-End-AAV-fwd/mCer-Ter-AAV-rev and mVen-End-AAV-fwd/mVen-Ter-AAV-rev. The four fragments were then assembled by Gibson Assembly and plasmids were verified by Sanger sequencing.

**Cloning of TopoI or TopoI mutants and construction of the pGB2-Para-TopoI plasmid**

pGB2-Para_TopoI was constructed in two steps using restriction cloning and Gibson-assembly. First, TopoI wildtype or mutants were PCR-amplified from genomic DNA extracted from *E. coli* MG1655 (PureLinkI®Genomic DNA Mini Kit, primers TopA-NheISD-fwd and TopA-SmaI-rev) and inserted into plasmid pBAD24 downstream of the arabinose-inducible promoter (Para) using restriction sites *Nhe*I and *Sma*I. Second, a fragment covering the gene araC, Para, the TopoI variant, and the *Amp*R resistance marker was PCR amplified (primers AraC-fwd, Amp-rev) from pBAD24 and fused with a PCR-amplified fragment (primers: pGB2-fwd, pGB2-rev) using the low-copy number plasmid pGB2 as template to obtain pGB2-Para-TopoI.

**TopoI editing in *E.coli* chromosome**

For TopoI degradation, *topA* was fused with the sequence encoding the PDT tag in the *mf*-lon strain described in[24] by Lambda-Red recombineering using pKD46 plasmid. Recombination cassettes were obtained by PCR from pECT3A plasmid with flanking regions homologous TopA-PDT-up/TopA-PDT-do. Excision of FRT-flanked Cm was mediated by transformation with the FLP recombinase-expressing plasmid pCP20.

Editing of the MG1655 *topA* gene to introduce the $Gly_5$, G703P or Y704A mutations was performed using the pCas/pTargetF system described in[52]. Twenty nucleotides were cloned in BsaI site in pEcgRNA, using primers gRNA-topA-1661_1681-fwd/gRNA-topA-1661_1681-rev for $Gly_5$, gRNA-topA-2444_2464-fwd/gRNA-topA-2444_2464-rev for G703P and Y704A. Gene editing was performed by Lambda-Red recombineering using pEcCas plasmid[52] and pEcgRNA-topA-2444_1681 or pEcRNA-topA-1661_1681. Recombination cassettes were obtained by PCR from pAX7-EcTopI-$Gly_5$, pAX7-EcTopI-G703P and pAX7-EcTopI-Y704A with TopA-mut-1661_1681-fwd/TopA-mut-1661_1681-rev for $Gly_5$, TopA-mut-2444_2464-fwd/TopA-mut-2444_2464-rev for G703P and Y704A. Mutation was verified by sanger sequencing. Plasmids were cured as described in[52].

#### Chloramphenicol gene insertion and *topA* mutations transduction

To transduce and select the $Gly_5$, G703P and Y704A mutations, the chloramphenicol gene was inserted between *btuR* and *rluB*, at 3.3 kb of *topA*, by Lambda-Red recombineering using pKD46 plasmid. Recombination cassettes were obtained by PCR from pKD3 plasmid with flanking regions homologous to *btuR* and *rluB* intergenic region using insert-Cm-3313-up/insert-Cm-3313-do. The chloramphenicol gene was inserted in 7 strains: MG1655 WT, TopoI-$Gly_5$ #1 and #2, TopoI-G703P #1 and #2 and TopoI-Y704A #1 and #2. These strains were used to generate P1 lysate, and further transduced into MG1655 WT.

#### Western blot

Bacteria were resuspended in Laemmli buffer at $2.00 \times 10^6$ cells/µL. Protein extracts were migrated on a 7.5 % gel and transferred to a membrane saturated with 10 % milk in TBS-T, then labelled with rabbit anti-TopA antibodies and finally revealed with horseradish peroxidase-coupled anti-rabbit antibodies. Revealing was carried out on the pico with the FX fusion device. Membranes were washed, and labelled with horseradish peroxidase-coupled anti-RpoB antibodies (a loading indicator of protein quantity). Revelation was performed at pico using the FX fusion device. For quantification, the grey level was calculated for each protein, the background was subtracted and the amount of TopoI was normalized to the amount of protein (loading control, RpoB).

#### One-dimensional gel electrophoresis with chloroquine

Plasmid DNA molecules were purified from bacteria in the exponential growth phase using a commercially available purification kit (Monarch® Plasmid Miniprep Kit, NEB). The purified plasmids (0.5-1 µg) were run on a 1.2 % agarose gel containing 2.5 µg/mL of chloroquine in 1x TBE buffer with 2.5 µg/mL of chloroquine at 25 V for 24 hours at 4 °C. The agarose gels were then washed in tap water three times during 1 hour, and stained by ethidium bromide.

#### RNA extraction and RNAseq

After 4 h induction of degradation, total RNA was extracted from *E.coli* WT and TopoI-PDT by an aqua-phenol extraction (ROTI®Aqua-Phénol) and ethanol precipitation. The RNA was then resuspended in diethyl pyrocarbonate-treated water. DNase treatment, rRNA depletion using

NEBNext rRNA Depletion kit and libraries were prepared with TruSeq Stranded Total RNA library Prep Gold kit and sequenced by Macrogen Europe as 101 pb paired end reads using Illumina Truseq. One biological replicate was generated for each condition, and an average ~80 million reads were generated per sample. Data were analysed with Rockhopper software.

#### Whole-genome sequencing and mutations identification

The genomic DNA was isolated from an exponential culture of *E. coli* MG1655 WT, TopoI-$Gly_5$ #1 and #2, TopoI-G703P #1 and #2 and TopoI-Y704A #1 and #2 using PureLinkI®Genomic DNA Mini Kit. NGS libraries for paired-end sequencing were prepared using Invitrogen Colibri PS DNA Library Prep Kit for Illumina according to the manufacturer's instructions. Raw reads were analysed directly in the Breseq pipeline to identify and annotate mutations compared to *E. coli* MG1655 (NC_000913.3).

#### Motility assay

Soft gel plates with 0.3 % agar in LB medium were prepared just before the start of the assay. A colony was picked into the centre of the plate. Plates were incubated at 30 °C for 18 hours. After incubation, the plates were imaged and the area of motility measured using ImageJ[53].

#### Gene expression assay

Glycerol stocks were streaked on resistance agar plates. Single colonies were inoculated in 1 mL of supplemented M9 minimal medium containing kanamycin at 50 µg/mL within a 2 mL 96 deepwell plate. Cultures were grown overnight in a thermoblock, at 37 °C and 300 rpm. The next day, cultures underwent a 1:500 dilution in supplemented M9 minimal medium with kanamycin, within a 2 mL 96 deepwell plate. Cultures were grown in a thermoblock, at 37 °C and 300 rpm until $OD_{600} \approx 0.1$. Cultures then underwent a 1:1000 dilution in supplemented M9 minimal medium with kanamycin. 200 µL of these diluted cultures were distributed in a 96 well plate (with black walls and a clear flat bottom), placed in a humidity cassette. Time series were acquired in a Tecan Spark Microplate Reader. Temperature was set at 37 °C. Continuous shaking was set on double orbital with an amplitude of 3 mm and a frequency of 90 rpm. Time points were acquired every 10 minutes, over a total period of 20 h. Three quantities were measured at each time point: absorbance (at 600 nm), mCerulean fluorescence (excitation at 430 nm and emission at 475 nm) and mVenus fluorescence (excitation at 510 nm and emission at 550 nm). The values of fluorescence measurements of each well over a period of 1.5 hours corresponding to late exponential phase were averaged and normalized by values of WT condition.

#### Nucleic acid sequences

| **Oligonucleotides for nucleic acid scaffolds* (5' → 3')** | |
|---|---|
| all scaffolds | RNA: GAGUCCGCGGCGCG |
| duplex scaffold | tDNA: CTCTGAATCTCTTCCGACGCGCCGCGGGACGTACTGACC |
| | ntDNA: GGTCAGTACGTCCTATCGATCTTCGGAAGAGATTCAGAG |
| minimal scaffold | tDNA: CTCTGAATCTCTTCCGACGCGCCGCG |
| | ntDNA: TCGGAAGAGATTCAGAG |
| short overhang scaffold | tDNA: CTCTGAATCTCTTCCGACGCGCCGCGGGACGTACTGACCTTAGAT |

| | |
|---|---|
| | ntDNA: TAGATTGGTCAGTACGTCCTATCGATCTTCGGAAGAGATTCAGAG |
| bubble scaffold | tDNA: CTCTGAATCTCTTCCGACGCGCCGCGAAAATTTTTTTTAAAATTATTAATCCC |
| | ntDNA: GGGATTAATAAttttttcattatttTTATCGATCTTCGGAAGAGATTCAGAG |
| long overhang scaffold | tDNA: CTCTGAATCTCTTCCGACGCGCCGCGAAAATTTTTTTTTTTTTTTTTTT |
| | ntDNA: TTAGTTAGtttttttcattatttTTATCGATCTTCGGAAGAGATTCAGAG |
| **Primers for TopoI mutants cloning** (5' → 3')** | |
| TopoI-G703P | Fw: CCGCATTAAAccgTATGACGGCCCGATCGTTGAGTGTG |
| | Rv: GGCCGTCATAcggTTTAATGCGGAATTCGCCCTCTTC |
| TopoI-Y704A | Fw: TTCCGCATTAAAGGTgcgGACGGCCCGATCGTTGAGTGTGAAAAATGTG |
| | Rv: CGATCGGGCCGTCcgcACCTTTAATGCGGAATTCGCCCTCTTCG |
| TopoI-Gly$_5$ | Fw: ggcggtggtggcgggATCTTACCAGCAGTTAATAAAGGCGATGCTC |
| | Rv: cccgccaccaccgccCAACGCAGGCATCACTTTTGTCCAG |
| **Oligonucleotides used for cloning, recombineering and plasmid construction (5' → 3')** | |
| gRNA-topA-2444_2464-fwd | TAGTTTCCGCATTAAAGGTTATGA |
| gRNA-topA-2444_2464-rev | AAACTCATAACCTTTAATGCGGAA |
| gRNA-topA-1661_1681-fwd | TAGTGTGATGCCTGCGTTGCGTAA |
| gRNA-topA-1661_1681-rev | AAACTTACGCAACGCAGGCATCAC |
| TopA-mut-1661_1681-fwd | GGCACGGCGATGGACAGC |
| TopA-mut-1661_1681-rev | CACTCTTCGTTGGTGCAGG |
| TopA-mut-2444_2464-fwd | GCCAGATGACCCCAGCG |
| TopA-mut-2444_2464-rev | CGGCTTGGTAAAGTGCTGG |
| mCer_Ter_fwd | CTCGAGACAATAACTGAATA |
| mVen_End_rev | TTTATACAGTTCGTCCATAC |
| mVen_Ter_fwd | AAGCTTAATTAGCTGAGTCT |
| mCer_End_rev | CTTATACAATTCATCCATGC |
| mCer_End_AAV_fwd | CCATGGCATGGATGAATTGTATAAGAGGCCTGCAGCAAACGAC |
| mCer_Ter_AAV_rev | CGCCCTATTCAGTTATTGTCTCGAGTTAAACTGCTGCAGCGTAGT |
| mVen_End_AAV_fwd | CCACGGTATGGACGAACTGTATAAAAGGCCTGCAGCAAACGAC |
| mVen_Ter_AAV_rev | CCTCTAGACTCAGCTAATTAAGCTTTTAAACTGCTGCAGCGTAGT |
| dusB-fwd-verif | GTTTGGCCTTTCATCTCGTGC |
| dusB-rev-verif | CGGTAGAAACGGTCAGTACG |
| cyoE-fwd-verif | CTGGTTGTAGGCTCCATCTGG |
| cyoE-rev-verif | CGAATGGGGTAAACATGCGG |
| insert-Cm-3313-up | TTTGATGCCGGTGTAAAAGCGCAGATAGGGATCGATTATTAATCTCTGACATGGGAATTAGCCATGG |
| insert-Cm-3313-do | AACGGTTAATCAAAGTATGCCGGACGTCATATCCGGCATTTTTACGTGTAGGCTGGAGCTGCTTC |
| TopA-PDT-up | TCAGCATTTTATGTTGATGGCAAATGGGTTGAAGGAAAAAAAGTTGGAGCGGCGAACAAA |
| TopA-PDT-down | CGCGACCCTTTGTTTATAAAAACCTGACAGAATTAAAGGTTAGTCGGATGTAGGCTGGAG |
| TopA-NheISD-fwd | AAAGCTAGCAGGAGGAATTCACCATGGGTAAAGCTCTTGTC |
| TopA-SmaI-rev | AAACCCGGGTTATTTTTTTCCTTCAACCCATTTG |

| | |
|---|---|
| pGB2-fwd | GATACCGCTGCCTTACTGGG |
| pGB2-rev | CGGATGAAGTGGTTCGCATCC |
| AraC-fwd | CCGAGGATGCGAACCACTTCATCCGCATAGGGTCATGGCTGCGC |
| Amp-rev | ATGCACCCAGTAAGGCAGCGGTATCCTACAGGGCGCGTAAATCAATC |

* The TopoI consensus cleavage sequence[15] is shown in lowercase.
** The mutated sequence is shown in lowercase.

### List of strains

| ***E.coli* strain name** | **Strain number** | **Genotype** | **Reference** |
|---|---|---|---|
| MG1655 | OE1 | K-12 MG1655 WT | Lab collection |
| TopoI-G703P #1 | OE3618 | OE1 *topA*-G703P clone 1 | This work |
| TopoI-G703P #2 | OE3619 | OE1 *topA* -G703P clone 2 | This work |
| TopoI-Y704A #1 | OE3628 | OE1 *topA* -Y704A clone 1 | This work |
| TopoI-Y704A #2 | OE3629 | OE1 *topA* -Y704A clone 2 | This work |
| TopoI-Gly5 #1 | OE3616 | OE1 *topA* -Gly5 clone 1 | This work |
| TopoI-Gly5 #2 | OE3617 | OE1 *topA* -Gly5 clone 2 | This work |
| *mf*-lon | OE3082 | MG1655pro *lacZ*::$P_{LtetO}$-*mf*-Lon | [24] |
| *mf*-lon TopoI-PDT | OE3437 | OE3082 *topA*-PDT3A | This work |
| LACRII | N/A | a derivative of *E. coli* LOBSTR protein expression strain with the two most abundant RNAses knocked out (*rna*-, *rnb*-) | [36,54] |

### List of plasmids

| **Plasmid name** | **Details** | **Reference** |
|---|---|---|
| LS | pSC101-derived plasmid, lacO-3205pb-$P_{01(apFAB67)}$-mCer-lacO-$P_{\lambda PR}$-mVen, KanR | [26] |
| pUC18 | *E.coli* cloning vector, $ori^T$, $Amp^R$ | Addgene |
| pUC19 | *E.coli* cloning vector, $ori^T$, $Amp^R$ | Addgene |
| pGB2-Para | pSC101-derived plasmid, *araC*, $P_{ara}$, $Amp^R$ | This work |
| pGB2-Para-TopoI-WT | pSC101-derived plasmid, *araC*, $P_{ara}$-*topA*, $Amp^R$ | This work |
| pGB2-Para-TopoI-Gly5 | pSC101-derived plasmid, *araC*, $P_{ara}$-*topA*-Gly5, $Amp^R$ | This work |
| pGB2-Para-TopoI-G703P | pSC101-derived plasmid, *araC*, $P_{ara}$- *topA*-G703P, $Amp^R$ | This work |
| pGB2-Para-TopoI-Y704A | pSC101-derived plasmid, *araC*, $P_{ara}$- *topA*-Y704A, $Amp^R$ | This work |
| pVS11_rpoA_rpoB_rpoC_HRV3C_His10_rpoZ | encoding all subunits of *E. coli* core RNAP enzyme with a $His_{10}$-tag on the CTD of β' subunit | [55] |
| pACYC_Duet1_rpoZ | encoding ω subunit of *E. coli* RNAP | [55] |
| pAX7-EcTopI | pET-derived plasmid, with a $His_{10}$-tag (cleavable with HRV3C protease) and a TwinStrep tag on the NTD of *E. coli* TopoI | This work |
